# Mitotic chromosomes scale to nucleo-cytoplasmic ratio and cell size in *Xenopus*

**DOI:** 10.1101/2022.10.20.513104

**Authors:** Coral Y. Zhou, Bastiaan Dekker, Ziyuan Liu, Hilda Cabrera, Joel Ryan, Job Dekker, Rebecca Heald

**Affiliations:** Department of Molecular and Cell Biology, University of California, Berkeley, Berkeley, CA, USA; Department of Systems Biology, University of Massachusetts Chan Medical School, Worchester, MA, USA; Advanced BioImaging Facility, McGill University, Montreal, Quebec, CA; Howard Hughes Medical Institute, Chevy Chase, MD, USA

## Abstract

During the rapid and reductive cleavage divisions of early embryogenesis, subcellular structures such as the nucleus and mitotic spindle scale to decreasing cell size. Mitotic chromosomes also decrease in size during development, presumably to coordinately scale with mitotic spindles, but underlying mechanisms are unclear. Here we combine *in vivo* and *in vitro* approaches using eggs and embryos from the frog *Xenopus laevis* to show that mitotic chromosome scaling is mechanistically distinct from other forms of subcellular scaling. We found that mitotic chromosomes scale continuously with cell, spindle and nuclear size *in vivo*. However, unlike for spindles and nuclei, mitotic chromosome size cannot be re-set by cytoplasmic factors from earlier developmental stages. *In vitro*, increasing nucleo-cytoplasmic (N/C) ratio is sufficient to recapitulate mitotic chromosome scaling, but not nuclear or spindle scaling, through differential loading of maternal factors during interphase. An additional pathway involving importin α scales mitotic chromosomes to cell surface area/volume (SA/V) during metaphase. Finally, single-chromosome immunofluorescence and analysis of Hi-C data suggest that mitotic chromosomes scale through decreased recruitment of condensin I, resulting in major rearrangements of DNA loop architecture to accommodate the same amount of DNA on a shorter axis. Together, our findings demonstrate how mitotic chromosome size is set by spatially and temporally distinct developmental cues in the early embryo.

## Introduction

Upon fertilization, embryos undergo a series of rapid cell division events in the absence of cell growth, thereby decreasing cell size. Subcellular structures including the nucleus and mitotic spindle scale to cell size through a set of defined mechanisms (Heald and Gibeaux, 2018; Levy and Heald, 2015). Mitotic chromosomes also shrink in size during development and scale with cell size across metazoans (Kramer et al., 2021; Micheli et al., 1993), but underlying mechanisms are poorly understood. In plants and in fly embryos, artificially lengthening chromosomes resulted in increased chromosome mis-segregation during mitosis (Schubert and Oud, 1997; Sullivan et al., 1993). Similar experiments in budding yeast showed that the artificially longer chromosome was hyper-compacted during anaphase due to Aurora B kinase phosphorylation of substrates including condensin, a key regulator of mitotic chromosome condensation and resolution (Neurohr et al., 2011). In *C. elegans*, a genetic screen for genes required for segregation of an extra-long chromosome identified the centromeric histone CENP-A and topoisomerase II (topo II) as regulators of holocentric chromosome size (Ladouceur et al., 2017). However, it is unclear whether pathways that tune the length of an artificially long chromosome also operate during the physiological process of mitotic chromosome scaling during embryogenesis.

Mechanisms that scale the spindle and nucleus during development have been well-characterized. As cell volume decreases, structural components become limiting (Good et al., 2013; Hazel et al., 2013). In addition, some scaling factors are regulated by the nuclear transport factor importin α, which partitions between the cytoplasm and the cell membrane and serves as a sensor for the cell surface area to volume ratio (Brownlee and Heald, 2019). Previous studies of mitotic chromosome scaling, performed mainly in *C. elegans*, revealed that mitotic chromosome size correlates positively with cell size and nuclear size and negatively with intranuclear DNA density (Hara et al., 2013; Ladouceur et al., 2015). Knockdown of importin α or the chromatin-bound Ran guanine exchange factor RCC1 decreased both nuclear and mitotic chromosome size (Hara et al., 2013; Ladouceur et al., 2015). Haploid embryos generated by katanin knockdown contained longer mitotic chromosomes compared to diploids (Hara et al., 2013). However, conserved relationships among genome size, nuclear size, and cell size complicate efforts to distinguish correlation from causation of mitotic chromosome scaling. Furthermore, it is unclear whether similar underlying mechanisms operate during embryogenesis of vertebrates that possess larger chromosomes and more complex karyotypes.

The African clawed frog *Xenopus laevis* provides a powerful system for studying mechanisms of mitotic chromosome scaling. Female frogs produce thousands of eggs that enable isolation of undiluted and cell cycle-synchronized cytoplasm in the form of egg extracts that reproduce many cellular processes *in vitro* including mitotic chromosome condensation (Maresca and Heald, 2006). In addition, fertilized eggs divide synchronously allowing extracts to be prepared from embryos at different stages of development. Our previous work demonstrated that egg extracts can recapitulate a decrease in chromosome size between early and late blastula stages of development *in vitro* (Kieserman and Heald, 2014), but did not uncover underlying scaling mechanisms. Here, we fully leverage the *Xenopus* system by systematically comparing changes in mitotic chromosome size observed *in vivo* with perturbations *in vitro* to distinguish factors that regulate mitotic chromosome scaling including nuclear size, spindle size, cell size, cell-cycle stage and nucleo-cytoplasmic (N/C) ratio. We find that mitotic chromosomes scale continuously with spindle size, even in large cells of the early embryo. We show that scaling occurs primarily through differential recruitment of the DNA loop extruding motor condensin I, which alters DNA loop size and thus length-wise compaction of chromosomes. Finally, we describe how two distinct developmental cues, cell size and nucleocytoplasmic ratio, combine to reduce chromosome length over the course of development. Together, these results create a multi-scale model for how mitotic chromosome size is set in the embryo and open up new avenues for deeper exploration of how changes in chromosome size and architecture contribute to early vertebrate embryogenesis.

## Results

### Mitotic chromosomes scale continuously with cell, nuclear, and spindle size

We reasoned that mitotic chromosome size may relate to nuclear size and content due to factors associated with the DNA prior to entry into mitosis, such as histones. Alternatively, mitotic chromosomes could scale with spindle size through mechanisms operating in mitosis. To distinguish between these possibilities, we performed a time course of whole embryo immunofluorescence through the late blastula stages of *X. laevis* development and measured the dimensions of cells, spindles, and metaphase plates (Figure 1A-B). Although previous work showed that spindle lengths reach a plateau in cells larger than ~200 microns in diameter (Figure 1-S1A, (Wühr et al., 2008)), measurement of spindle volumes by confocal microscopy revealed size scaling in cells as large as 600 microns in diameter (Figure 1C), consistent with observations that spindle width correlates more robustly with cell volume than spindle length in cultured cells (Figure 1-S1B, (Kletter et al., 2021). Mitotic chromosome volumes also scaled continuously with cell size, similar to published work describing nuclear scaling (Figure 1D, Figure 1-S2, (Jevtić and Levy, 2015)). To assess whether mitotic chromosomes scale more with nuclear size or with mitotic spindle size, we binned the data by cell size and plotted average volumes of the different subcellular structures. We found that mitotic chromosomes scaled remarkably well with both spindles and nuclei (Figure 1-S3). However, the magnitude change in nuclear volume was far greater than for mitotic structures: nuclei scaled by ~10-fold over early cleavage divisions, while mitotic chromosomes and spindles scaled by 3-fold and 2-fold, respectively (Figure 1E-F). Thus, the change in chromosome compaction as cells enter mitosis diminishes from 8-fold in early blastula embryos to 1.5-fold in late blastula stages (Figure 1-S4). Overall, these results demonstrate that mitotic chromosomes scale continuously with cell size in the early embryo and suggest that mitotic chromosomes may share scaling features with both nuclei and mitotic spindles.

**Figure 1:**
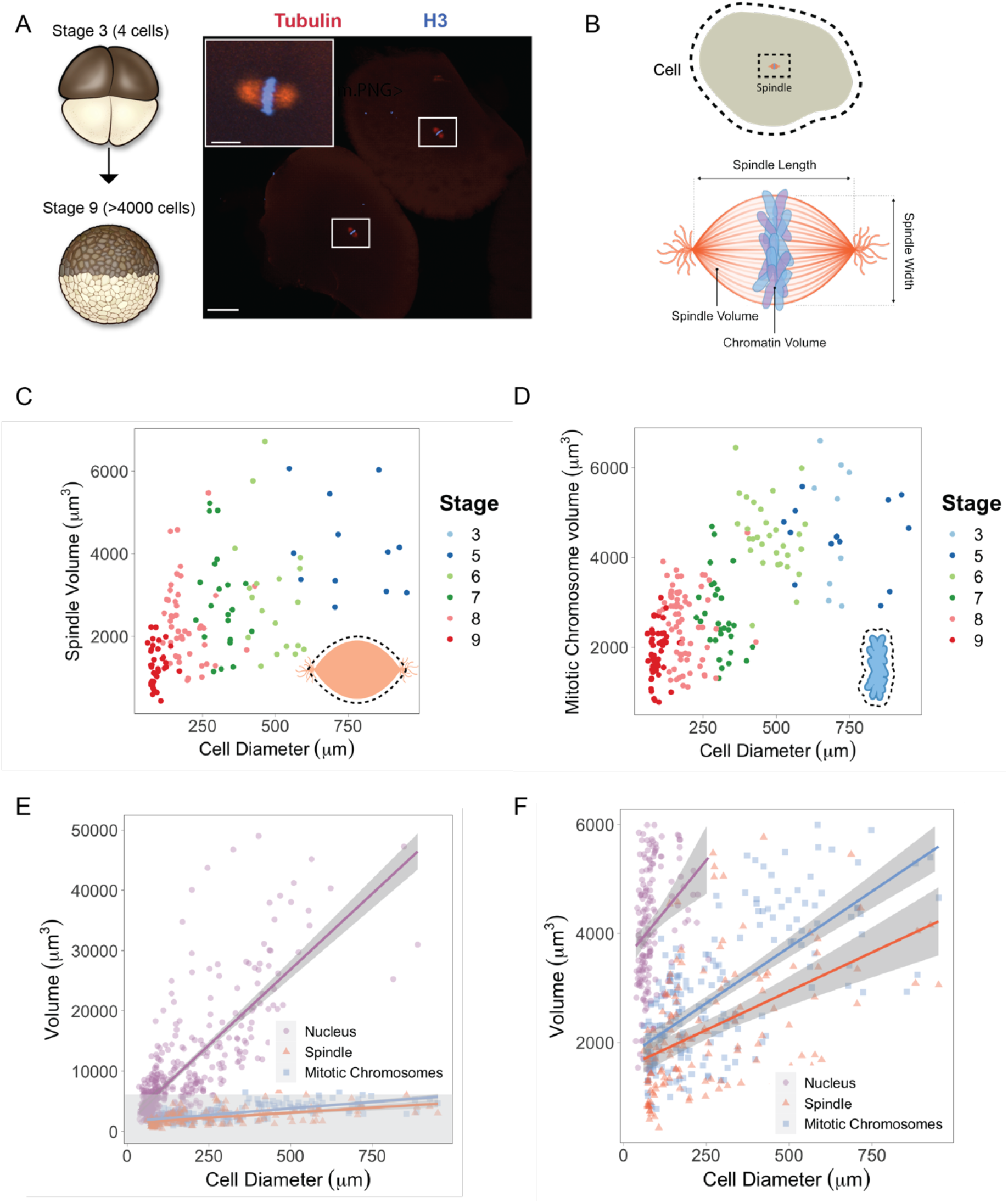
Mitotic chromosomes scale continuously with cell size. (A) Experimental scheme for whole-embryo immunofluorescence. Blastula-stage embryos were fixed during mitosis and stained with anti-histone H3 and anti-tubulin antibodies to visualize mitotic chromosomes and spindles, respectively. Representative image of two cells from a stage 6 embryo with white rectangles outlining mitotic spindles, scale bar = 100 μm. Inset: Magnified view of one of the mitotic spindles, scale bar = 20 μm. (B) Dimensions of cells and spindles were either directly measured or calculated (for details see Materials and Methods). (C) Measurements of spindle volume or (D) mitotic chromosome volume plotted against cell diameter, colored by developmental stage. (E) Volumes of spindles, nuclei and mitotic chromosomes all plotted against cell diameter, fit with linear models. 95% confidence intervals shown in gray. (F) Zoom-in of gray panel shown in (E). n = 2 biological replicates.

### Mitotic chromosomes scale length-wise

To examine how morphologies of individual mitotic chromosomes change during development, we prepared mitotic cell extracts from stage 3 (4-cell) or stage 8 (~4000-cell) embryos (Wilbur and Heald, 2013) and centrifuged single endogenous mitotic chromosomes onto coverslips for size measurements. We found that median chromosome lengths decreased ~2-fold between stage 3 and stage 8 (Figure 2A-B), similar to the magnitude of mitotic chromosome scaling during this period estimated by whole-embryo immunofluorescence (~2-3-fold, Figure 1D) and indicating that shortening of the long axis is the predominant metric underlying mitotic chromosome scaling during early embryogenesis. We also observed that the median length of endogenous stage 3 mitotic chromosomes was not statistically different from that of replicated sperm chromosomes formed in egg extracts (Figure 2B), demonstrating that replicated sperm chromosomes formed in egg extracts serve as a proxy for mitotic chromosome size during the earliest cell divisions.

**Figure 2:**
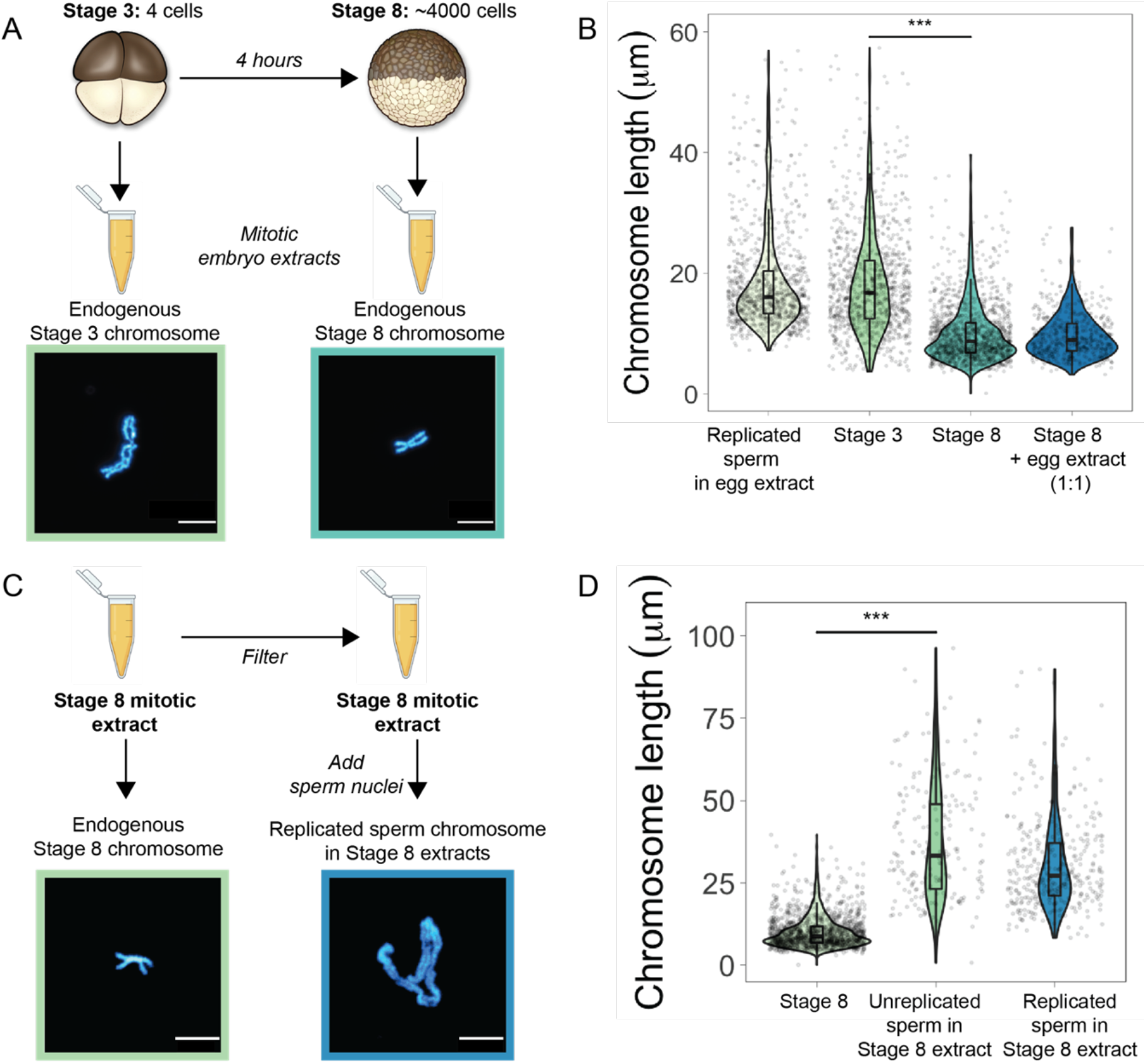
Mitotic chromosomes scale length-wise. (A) Mitotic extracts were prepared from stage 3 or stage 8 embryos and endogenous, individual mitotic chromosomes were centrifuged onto coverslips and stained with Hoechst DNA dye. Representative images of stage 3 and stage 8 chromosomes are shown. (B) Length distributions of sperm mitotic chromosomes replicated in egg extract, mitotic chromosomes isolated from embryo extracts and stage 8 embryo extract chromosomes after mixing 1:1 with egg extract. (C) Stage 8 extracts were filtered to remove endogenous chromosomes, then unreplicated or replicated sperm nuclei were added to form mitotic chromosomes. Representative images of endogenous stage 8 chromosome or replicated sperm chromosome formed in stage 8 extracts shown here. (D) Quantification of chromosome lengths for the experiment shown in (C). n=3 biological replicates, >50 chromosomes per replicate. Scale bar = 10 μm. *** denotes p <0.001 by one-way ANOVA statistical testing.

Previously, it was shown that mixing mitotic extracts prepared from early and late blastula stage embryos resulted in spindles of intermediate size due to equilibration of cytoplasmic spindle scaling factors (Wilbur and Heald, 2013). Likewise, combining interphase extracts at different ratios from two *Xenopus* species with different sized nuclei produced a graded effect on nuclear size (Levy and Heald, 2010). To test whether a similar mechanism operates on mitotic chromosomes, we combined metaphase-arrested egg extracts in a 1:1 ratio with stage 8 mitotic embryo extracts containing endogenous mitotic chromosomes (Figure 2B). However, we observed no increase in chromosome length, indicating that mitotic chromosome scaling factors are not exchangeable in the cytoplasm during metaphase. To test whether mitotic chromosome scaling could occur if initiated before the onset of chromosome condensation, we filtered stage 8 extracts to remove endogenous chromosomes and added back unreplicated sperm chromosomes or sperm nuclei that had undergone replication in egg extracts (Figure 2C). In both cases, sperm chromosomes were at least 2-fold longer than the endogenous stage 8 chromosomes (Figure 2D). Thus, mitotic chromosome size is predominantly set by factors loaded during interphase that are not exchangeable, making mitotic chromosome scaling fundamentally distinct from nuclear or spindle size scaling.

### Mitotic chromosome size is determined by nuclear factors during interphase

The results above indicated that mitotic chromosome size is largely determined by nuclear rather than cytoplasmic factors. Consistent with this idea, we confirmed previous results that G2-arrested nuclei from blastula-stage embryos added to metaphase egg extracts produced mitotic chromosomes ~2-fold shorter than replicated sperm chromosomes formed in the same extract (Figure 3A, C; (Kieserman and Heald, 2014)). This finding suggested that mitotic chromosome size is determined prior to entry into metaphase, likely by chromatin factors loaded during interphase. The 2-fold difference in chromosome size was also recapitulated in extracts depleted of membranes through ultracentrifugation, which are unable to form spindles but are competent for mitotic chromosome assembly, indicating that spindle formation is not required for mitotic chromosome scaling (Figure 3-S1). Previous work in *C. elegans* suggested that mitotic chromosome size correlates with intranuclear density and nuclear size (Hara et al., 2013), but we observed that embryo nuclei were larger than interphase sperm nuclei, and mitotic spindles formed in egg extracts with these two sources of nuclei were indistinguishable in size (Figure 3-S2). These data further suggest that scaling of spindles, nuclei and mitotic spindles are not necessarily coordinated.

**Figure 3:**
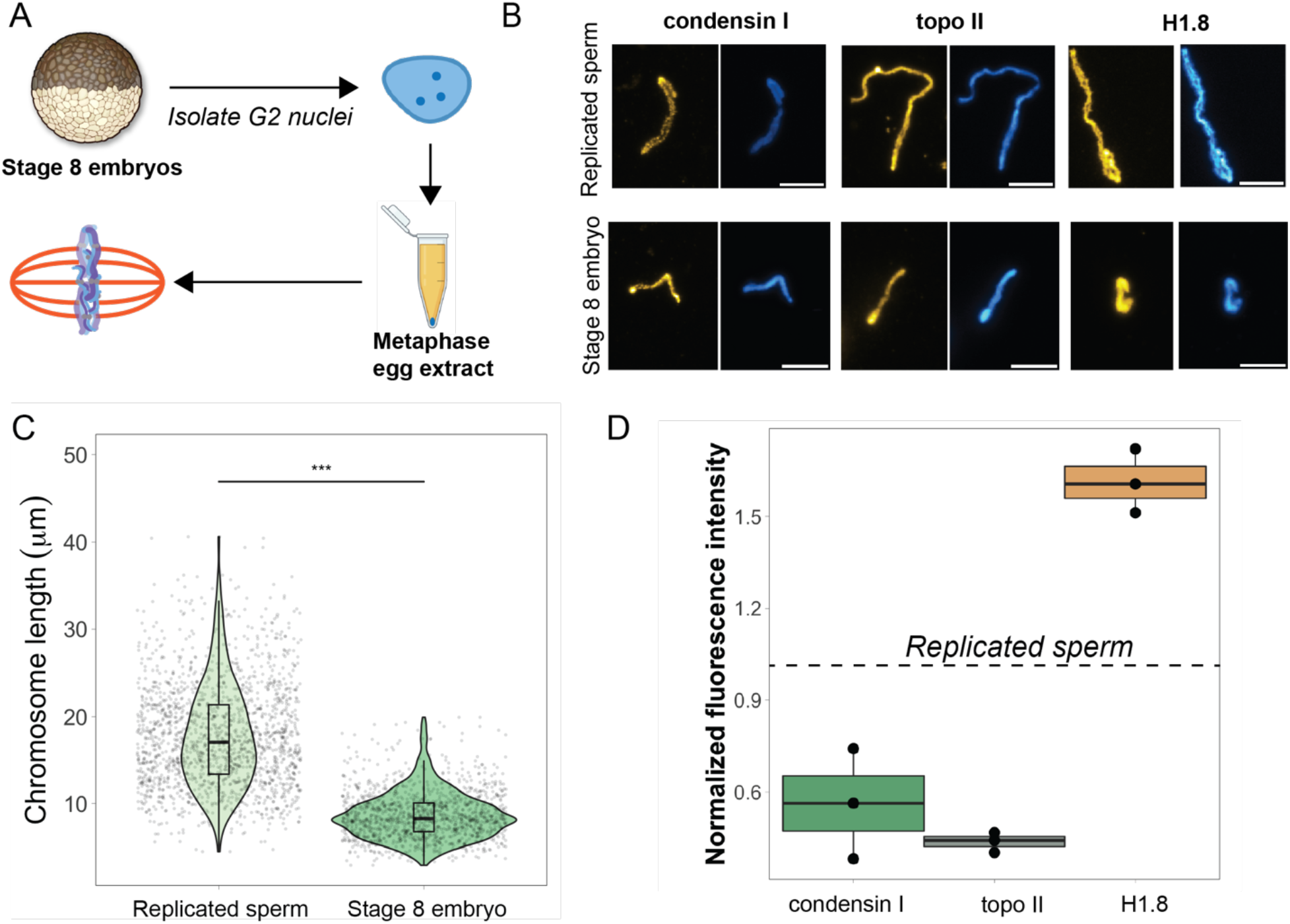
Egg extracts recapitulate mitotic chromosome scaling through differential recruitment of condensin I, topo II, and histone H1.8. (A) Experimental scheme. Stage 8 embryos arrested in G2 with cycloheximide were used to prepare extracts, then embryo nuclei were pelleted and added to metaphase-arrested egg extracts to form mitotic spindles and chromosomes. (B) Representative images of mitotic chromosomes prepared by adding replicated sperm nuclei (top) or stage 8 embryo nuclei (bottom) to metaphase egg extracts, and stained with antibodies for condensin I, topo II or H1.8. Scale bar = 10 μm. (C) Lengths of replicated sperm chromosomes or stage 8 embryo chromosomes formed in metaphase egg extracts. n>3 biological replicates, >50 chromosomes per replicate. *** denotes p <0.001 by one-way ANOVA statistical testing. (D) Abundances of topo II, condensin I and H1.8 (calculated by normalizing immunofluorescence signal to Hoechst signal, see Materials and Methods for details) on short embryo chromosomes normalized to long sperm chromosomes (shown by dotted line), from three different extracts.

Interestingly, although mitotic chromosome scaling could be recapitulated by adding embryo nuclei to metaphase-arrested egg extracts, chromosome morphologies were distinct. The separation of sister chromatid arms resulting in X-shaped mitotic chromosomes in both stage 3 and stage 8 mitotic embryo extracts (Figure 2A) was not observed when stage 8 embryo nuclei were added to egg extracts, as chromosome arms remained tightly associated along their lengths (Figure 3B). Taken together, these results indicate that factors determining mitotic chromosome size remain associated with G2-arrested embryo nuclei when placed into metaphase egg extracts, while factors required for regulating sister chromatid arm cohesion do not.

### Chromosome scaling correlates with differential recruitment of condensin I, topo II, and histone H1.8

Robust recapitulation of chromosome scaling in metaphase-arrested egg extracts enabled molecular-level analysis of potential scaling factors, which was not technically feasible in embryo extracts that cannot transit the cell cycle *in vitro*. We examined three proteins known to influence chromosome size and morphology in *Xenopus*, condensin I (the predominant condensin in *Xenopus* eggs), topoisomerase II (topo II), and the maternal linker histone, termed H1.8 (Maresca et al., 2005; Nielsen et al., 2020; Shintomi and Hirano, 2011). After performing immunostaining of short embryo chromosomes or long sperm chromosomes formed in the same egg extracts, the abundance of each factor was calculated by normalizing immunofluorescence signal to DNA dye intensity (Figure 3B, see Materials and Methods). We found that short embryo chromosomes contained less condensin I and topo II, but more H1.8 relative to long replicated sperm chromosomes (Figure 3D). These results are consistent with studies showing that depletion of H1.8 from egg extracts lengthens mitotic chromosomes (Maresca et al., 2005). Furthermore, it was recently shown that H1.8 inhibits binding of condensin I and topo II to mitotic chromosomes (Choppakatla et al., 2021). Therefore, differential recruitment of condensin, topo II, or H1.8 may contribute to mitotic chromosome scaling during embryogenesis.

Our previous work showed that short embryo chromosomes could be reset to lengths observed in replicated sperm samples by cycling the mitotic chromosomes through an additional interphase in egg extracts (Figure 3-S3A; (Kieserman and Heald, 2014)). To test whether the abundance of candidate scaling factors was affected, we performed immunofluorescence on mitotic embryo chromosomes before and after the additional interphase. We found that the abundance of all three factors on mitotic chromosomes increased on the embryo chromosomes (Figure 3-SC-F), which also doubled in length (Figure 3-S3B). Of the three factors, condensin I levels increased the most (2-fold), returning to levels found on replicated sperm chromosomes (Figure 3-S3C). Our observation that H1.8 levels increased slightly after the second metaphase suggests that condensin I abundance is not necessarily regulated by H1.8, and that condensin I can override the condensation activity of H1 to lengthen embryo chromosomes.

### Mitotic chromosomes scale through extensive remodeling of DNA loop architecture

Condensin shapes mitotic chromosomes through its ability to form and extrude loops from the central axis (Ganji et al., 2018; Goloborodko et al., 2016). *In silico* models of loop extrusion activity suggested that tuning the abundance of condensin could dramatically alter DNA loop architecture and thus chromosome dimensions (Goloborodko et al., 2016). However, these models have not been tested under physiological conditions that relate to chromosome size changes *in vivo*.

To assess how DNA loop size and arrangement is altered in the context of mitotic chromosome scaling, we performed Hi-C on long sperm chromosomes and short embryo chromosomes formed in egg extracts. Hi-C contact maps indicate that short embryo chromosomes had increased genomic contacts along their entire length, as evidenced by thickening of the diagonal (Figure 4A). To quantify this effect, we plotted the decay of contact frequencies *(P)* as a function of genomic distance in bp (*s*) (Figure 4B). We find that the shape of *P*(*s*) is similar to what we observed in earlier work for mitotic chromosomes from human, chicken and *Xenopus* (Choppakatla et al., 2021; Gibcus et al., 2018; Naumova et al., 2013), and for rod-shaped dinoflagellate chromosomes (Nand et al., 2021). *P*(*s*) is characterized by two regimes: an initial regime where the slope is rather small, followed by a regime where the slope is much steeper. We have shown that this shape is characteristic of compact rod-shaped chromosomes that are organized in layers (Naumova et al., 2013), and the genomic distance where the slope suddenly increases is related to the amount of DNA that is packed per layer. Loci within a layer often interact, but loci separated by a genomic distance that is larger than the layer size rarely do because of the stiffness of the chromosome. Within a layer, chromosomes are organized as loops, the size of which can be estimated from the derivative of *P*(*s*) (Johan H. Gibcus et al., 2018), the loop size is around where the derivative displays a maximum (Figure 4C).

**Figure 4:**
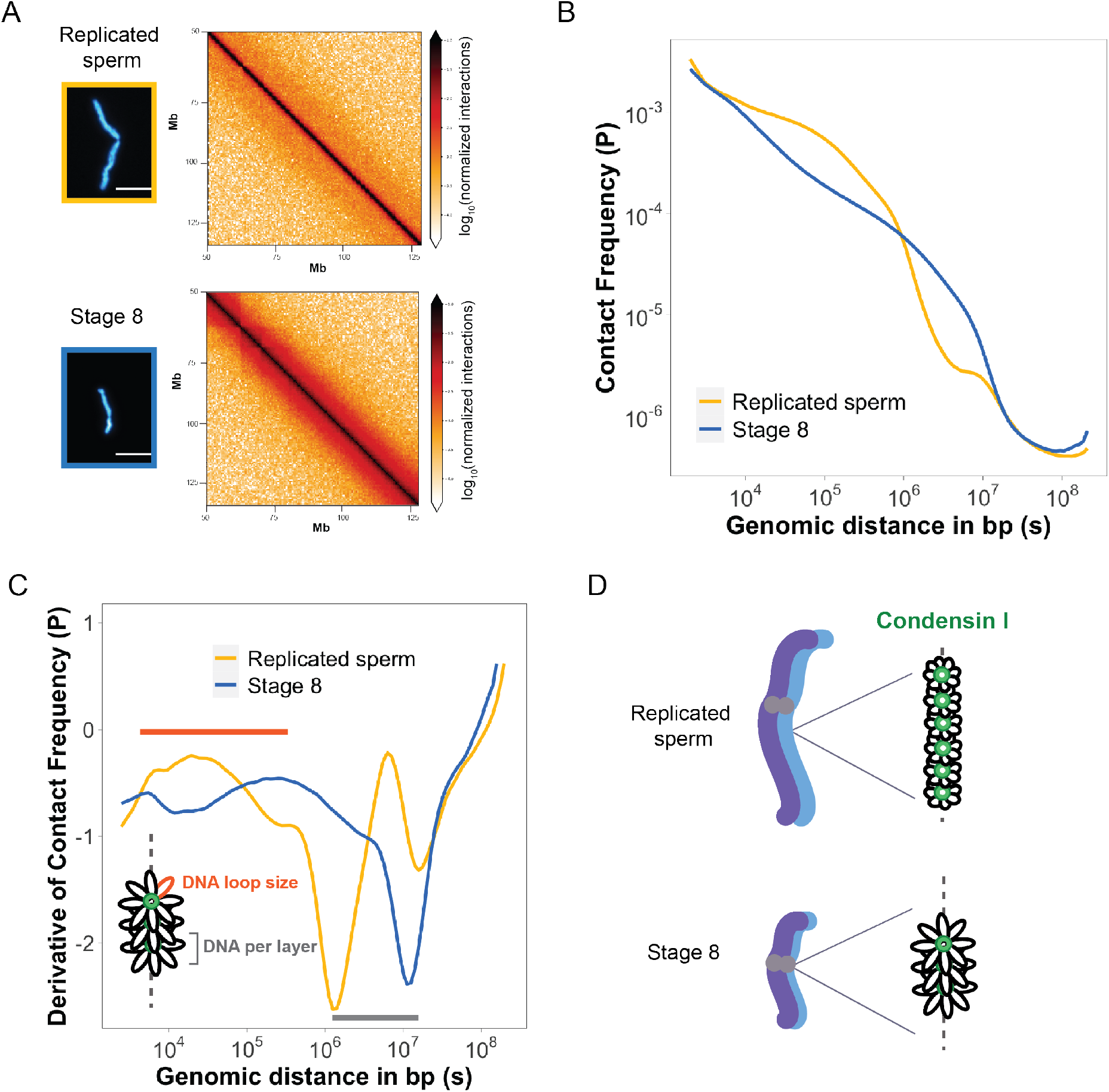
Mitotic chromosomes scale through extensive remodeling of DNA loop architecture. (A) Hi-C maps of chromosome 4S from replicated sperm or stage 8 embryo chromosomes formed in metaphase egg extracts. (B) Plots comparing how contact frequency *(P)* genome-wide decays as a function of genomic distance *(s)* for replicated sperm (yellow) or stage 8 mitotic chromosomes (blue). (C) Derivative of contact frequencies from (B). Based on previous models, the difference in inflection point at 10^4^-10^6^ bp represents a difference in loop size (orange), while the difference in inflection point at 10^6^-10^7^ bp represents a difference in DNA per layer (gray). (D) Model depicting how lower condensin I occupancy on short embryo chromosomes results in an increase in DNA loop size and DNA per layer. n= 2 biological replicates.

We find that *P*(*s*) for short embryo chromosomes differ in two ways from long sperm chromosomes. First the layer size is larger, i.e., ~10^7^ bp vs. 10^6^ bp (Figure 4C, gray bar), which would be expected for shorter chromosomes where more DNA is packed within a cross-section of a chromosome. Second, DNA loop size is considerably larger for short embryo chromosomes, as is visible in a rightward shift in the peak of the derivative of *P*(*s*) (Figure 4C, orange bar). This analysis, combined with our immunofluorescence results from Figure 3, is consistent with a model where mitotic chromosomes scale through decreased recruitment of condensin I, resulting in larger DNA loops and more DNA per layer, thus accommodating more DNA on a shorter chromosome axis (Figure 4D).

### Nucleo-cytoplasmic ratio regulates mitotic chromosome scaling, but not nuclear or spindle scaling

We next investigated the developmental cues that could decrease the abundance of condensin I on mitotic chromosomes as they scale during development. Characteristic features of cleavage divisions during early embryogenesis are the lack of cell growth and minimal gene expression, resulting in exponentially increasing copies of the genome within the same total volume and content of cytoplasm. The increase in nucleo-cytoplasmic (N/C) ratio, defined here as the number of nuclei per volume of cytoplasm, titrates a finite maternal pool of DNA binding factors that are distributed to more and more genomes over time, thus lowering their abundance per genome. This effect is thought to underlie activation of zygotic transcription at the mid-blastula transition (Amodeo et al., 2015), and titration of the histone chaperone Npm2 was shown to play a role in nuclear scaling (Chen et al., 2019).

To test whether N/C ratio could play a role in mitotic chromosome scaling, we varied the concentration of sperm nuclei at two different densities corresponding to early (~75 sperm nuclei/μl) and late (~1250 nuclei/μl) blastula stage embryos (Figure 5A). We first allowed these nuclei to replicate in interphase egg extracts, then added back fresh metaphase extracts before analyzing mitotic chromosome lengths and abundances of condensin I, topo II and H1.8. We found that increasing N/C ratio decreased mitotic chromosome size by ~1.5 fold (Figure 5B), consistent with the difference observed *in vivo* at the stages of development that correspond to the N/C ratios tested (Stage 6-7, Figure 1D). This 1.5-fold change in chromosome size also correlated with a 1.5-fold decrease in condensin I abundance on mitotic chromosomes (Figure 5C), and with less significant changes for H1.8 and topo II abundance (Figure 5-S1). Interestingly we found that increasing N/C ratio did not significantly affect nuclear size and actually led to an increase in spindle size (Figure 5-S2), suggesting that N/C ratio is only sufficient to scale mitotic chromosome size during embryogenesis.

**Figure 5:**
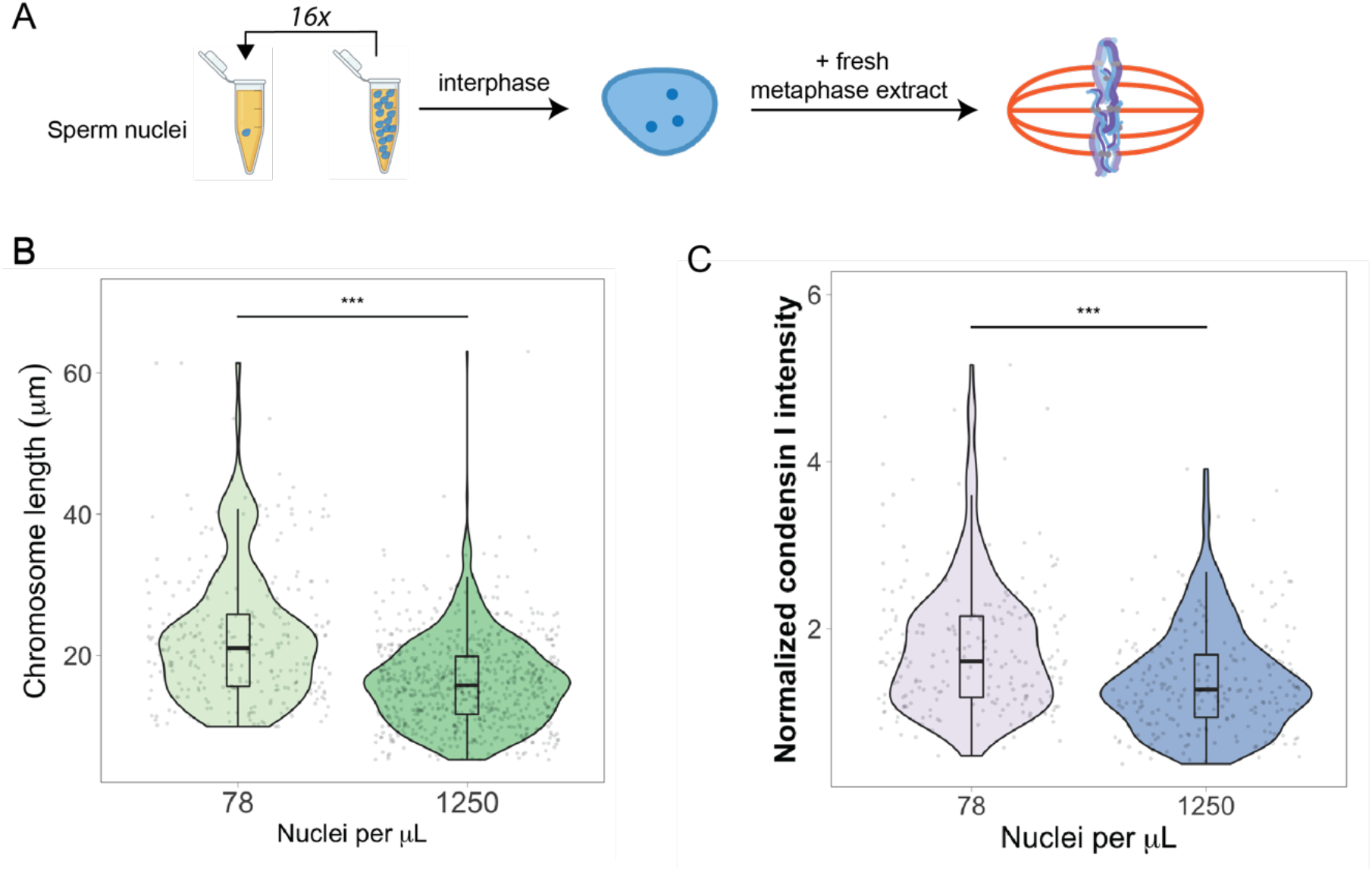
Nucleo-cytoplasmic ratio regulates mitotic chromosome scaling, but not nuclear or spindle scaling. (A) Concentration of sperm nuclei in egg extracts was varied 16-fold to mimic concentrations found in early (78 nuclei/μL) vs. late (1250 nuclei/μL) blastula stages. (B) Quantification of mitotic chromosome lengths and (C) condensin I intensities, normalized to Hoechst signal, in samples containing high or low concentrations of sperm nuclei. n=3 biological replicates, >50 chromosomes per replicate, *** denotes p <0.001 by one-way ANOVA statistical testing.

To examine whether the titration effect of chromosome factors observed *in vitro* corresponded to changes in their abundance during development *in vivo*, we measured levels of condensin I, H3 and histone H1.8 on embryo nuclei isolated from different stages. As predicted, although protein concentrations in embryos did not change over the course of the early cleavage divisions (Figure 5-S3A), levels of all factors on interphase nuclei decreased as genome copy number increased (Figure 5-S3B). Furthermore, adding different concentrations of stage 8 embryo nuclei to egg extracts did not significantly change mitotic chromosome size (Figure 5-S4), consistent with the idea that titration of maternal factors had already occurred in the embryo, and that interphase factors set chromosome size during metaphase. Together these observations confirm that maternally loaded factors are titrated onto newly synthesized copies of the genome, and that increasing N/C ratio is sufficient to shorten mitotic chromosomes, likely by decreasing levels of condensin I.

### Importin α partitioning scales mitotic chromosomes to spindle and cell size

A major developmental cue central to the scaling of nuclei and spindles is cell size. We previously identified importin α as a factor that coordinately scales nuclei and spindles to cell size by regulating nuclear import of lamin proteins and the activity of a microtubule destabilizing protein, respectively (Brownlee and Heald, 2019; Levy and Heald, 2010; Wilbur and Heald, 2013). Palmitoylation of importin α drives a subset of the total population to the cell membrane, where it can no longer interact with its nuclear localization sequence (NLS)-containing cargo that scale mitotic spindles (Figure 6A). As cell size decreases during early cleavage divisions, cell surface area/volume (SA/V) increases, causing more importin α to be driven to the membrane and more scaling factors to be released into the cytoplasm (Figure 6A; (Brownlee and Heald, 2019)). To address whether importin α also plays a role in mitotic chromosome scaling, we treated egg extracts with palmostatin, an inhibitor of the major depalmitoylation enzyme APT1 to increase the pool of palmitoylated importin α, thus mimicking smaller cells with higher cell SA/V (Figure 6B; (Dekker et al., 2010)). We found that mitotic chromosome size decreased with palmostatin treatment and was fully rescued by the addition of recombinantly purified importin α that cannot be palmitoylatated (NP importin α), but not by addition of wild-type (WT) importin α (Figure 6B-C). Surprisingly, we found no significant difference in condensin I abundance on mitotic chromosomes formed in either DMSO or palmostatin-treated extracts (Figure 6D), suggesting that importin α partitioning acts separately from the N/C pathway to scale mitotic chromosomes. To determine when in the cell cycle palmostatin was affecting importin α cargos, we added it to extracts either before interphase or following entry into metaphase (Figure 6E). Interestingly, analogous to effects on spindle size but not nuclear size (Brownlee and Heald, 2019), we found that palmostatin treatment during metaphase was sufficient to scale mitotic chromosomes (Figure 6F). Together these results demonstrate that partitioning of importin α during metaphase scales mitotic chromosomes to spindle and cell size in a condensin I-independent pathway.

**Figure 6:**
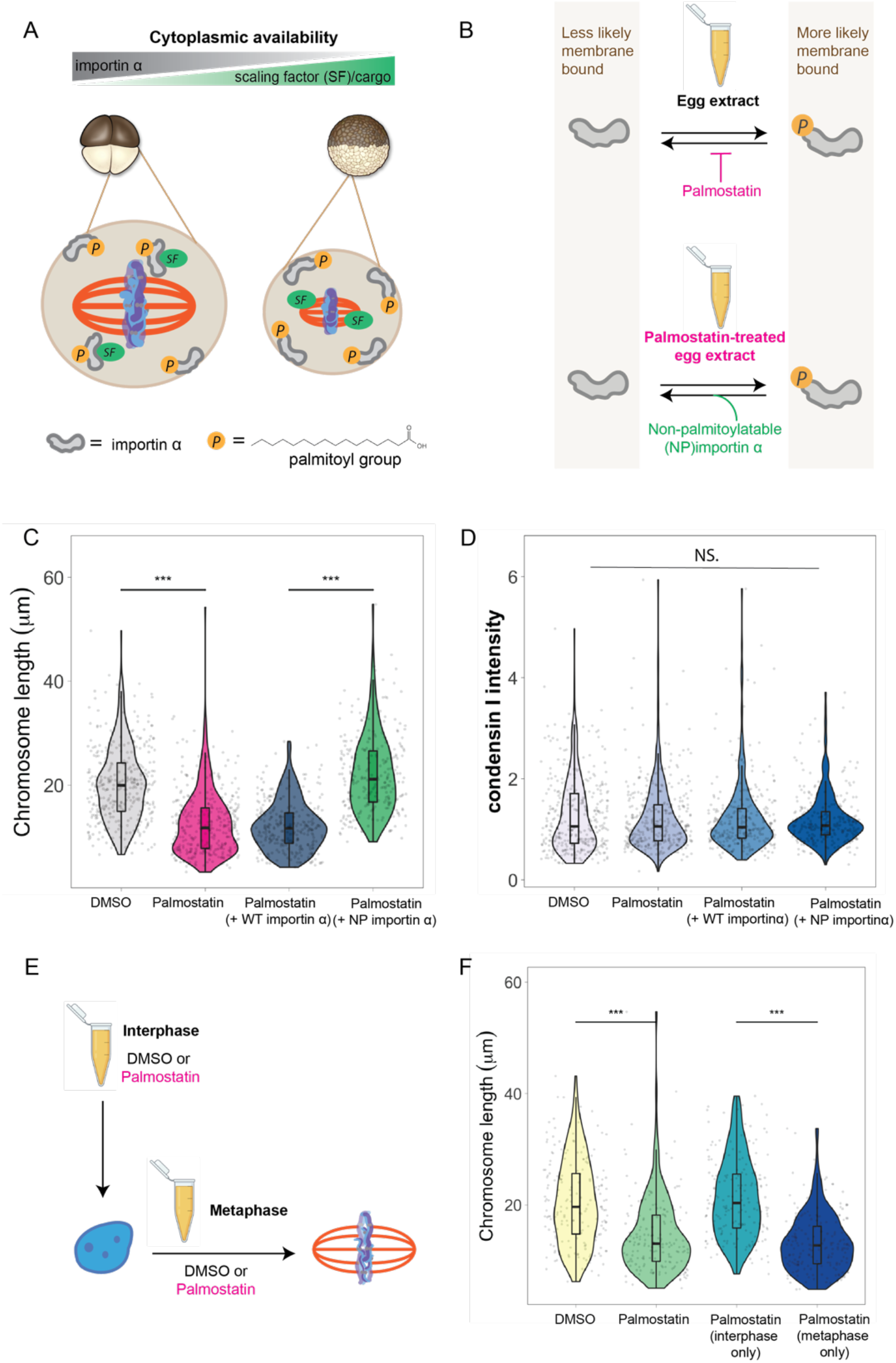
Importin α partitioning scales mitotic chromosomes to spindle and cell size. (A) Model for how importin α scales mitotic spindles to cell size. Due to palmitoylation of importin α, a portion of it is driven to the cell membrane, where it can no longer interact with nuclear localization sequence (NLS)-containing scaling factors, freeing them to shrink the mitotic spindle. As cell surface area/volume (SA/V) increases during embryogenesis, proportionally more importin α is driven to the membrane, thus increasing the cytoplasmic availability of scaling factors. (B) Top, Inhibition of the major depalmitoylation enzyme APT1 in egg extracts mimics smaller cells by increasing the proportion of palmitoylated, membrane-bound importin α. Bottom, addition of non-palmitoylatable (NP) importin α should rescue chromosome size in palmostatin-treated egg extracts by increasing the proportion of cytoplasmic importin α. (C) Quantification of mitotic chromosome lengths in palmostatin-treated extracts and rescue of chromosome length by addition of non-palmitoylatable (NP) importin α but not by wild-type (WT) importin α. (D) Quantification of condensin I intensity for the samples shown in (C). (E) Schematic of experiment to test whether importin α partitioning plays a role in chromosome scaling during interphase or metaphase. (F) Quantification of chromosome lengths for experiment described in (E). n=3 biological replicates, >50 chromosomes per replicate, and *** denotes p <0.001 by one-way ANOVA statistical testing.

## Discussion

The results described here provide a comprehensive view of how mitotic chromosome size is regulated by different developmental cues during early cleavage divisions of *Xenopus* embryos. Our data suggest that mitotic chromosomes coordinately scale with spindle and cell size through importin α partitioning. Additionally, increasing N/C ratio is sufficient to scale mitotic chromosomes through decreased recruitment of condensin I, resulting in increased DNA loop and layer size and length-wise compaction. As detailed below, our findings have important implications for the interplay between subcellular scaling and chromosome structure during early embryogenesis.

### Mitotic chromosome size is not necessarily coupled to nuclear and spindle size

Previous work in *C. elegans* showed that mitotic chromosome size correlated positively with nuclear size and negatively with intranuclear density (Hara et al., 2013; Ladouceur et al., 2015). However, in egg extracts, mitotic chromosome size does not necessarily correlate with spindle and nuclear size. G2-arrested stage 8 embryo nuclei, which contain both the maternal and paternal genomes, are almost 2-fold larger than replicated sperm nuclei (Figure 3-S2A), consistent with a 2-fold increase in genome content. Yet when added to metaphase egg extracts, they form mitotic chromosomes that are 2-fold shorter than replicated sperm chromosomes (Figure 3C), demonstrating that mitotic chromosomes don’t necessarily scale to either intranuclear density or nuclear size. We also showed that though mitotic chromosomes scale continuously with spindle size *in vivo* (Figure 1), this correlation is abolished *in vitro* (Figure 3-S2B). In a completely different set of experiments, when N/C ratio was varied in egg extracts (Figure 5), mitotic chromosomes shrank 1.5-fold while spindle size increased almost 2-fold and nuclear size remained constant. Together these data suggest that the mechanisms that regulate the size of mitotic chromosomes, spindles and nuclei are distinct.

### Mitotic chromosome scaling involves temporally and spatially distinct developmental cues

Whereas mitotic spindle size and nuclear size are set by factors operating during metaphase and interphase, respectively, we found that both phases of the cell cycle contribute to scale mitotic chromosomes. Our embryo extract and egg extract data (Figures 2 and 3) suggest that mitotic chromosome size is determined by factors already present in interphase nuclei and cannot be re-set by metaphase egg extracts. On the other hand, importin α partitioning plays an additional role to scale mitotic chromosomes to spindle and cell size specifically during metaphase. Consistent with this temporal separation of developmental cues, condensin I acts as a scaling factor only in the N/C pathway and not the importin α pathway (Figure 5C vs. Figure 6D). Together, these data suggest that the NLS-containing cargo that scale mitotic chromosomes, unlike cargo that scale nuclei and spindles, cannot freely exchange with other factors in the egg cytoplasm to lengthen short embryo chromosomes. Future work will be required to identify this cargo. Finally, since *Xenopus* embryos divide asymmetrically, with larger cells on the vegetal side, our results suggest that within the same embryo, the importin α and N/C ratio pathways could combine to have different effects on mitotic chromosome size.

### Multiple molecular pathways can regulate mitotic chromosome size

Based on *in silico* models of condensin I loop extrusion activity, it was predicted that changing condensin I occupancy on mitotic chromosomes causes major changes in DNA loop size and chromosome dimensions (Goloborodko et al., 2016). However, it was unclear whether such changes occur *in vivo*. Our results suggest that within the physiologically relevant range of condensin I concentrations present during embryogenesis, less condensin I correlates with both increased loop size and increased length-wise compaction. Another recent study showed that H1.8 can suppress condensin I occupancy on sperm mitotic chromosomes, reducing their length (Choppakatla et al., 2021). In contrast, we found that an increase in condensin I could act independently of H1.8 to lengthen chromosomes (Figure 3-S3), suggesting that condensin I is the major scaling factor for mitotic chromosomes. In the future it will be interesting to examine other factors that could be acting upstream or downstream of condensin I to set mitotic chromosome size in the embryo.

### N/C ratio as a fundamental mechanism for regulating chromatin structure during pre-ZGA cleavage divisions

Previous work in zebrafish and frogs suggested that titration of maternal factors such as histone H3 due to increasing N/C ratio plays an important role in regulating timing of Zygotic Genome Activation (ZGA) in the embryo (Amodeo et al., 2015; Joseph et al., 2017). One proposed model is that decreasing histone abundance facilitates the binding of transcription activation machinery (Joseph et al., 2017). Here we were able to recapitulate this titration effect by simply increasing the concentration of nuclei in egg extracts (Figure 5) and found that N/C ratio is sufficient to regulate mitotic chromosome size but not spindle or nuclear size. Together these results suggest that N/C ratio could be a universal mechanism for regulating chromatin structure across the cell cycle, but for completely different functions: transcriptional regulation during interphase and chromosome segregation during metaphase. In the future it will be interesting to identify additional factors that are titrated from maternal cytoplasm, and how this affects overall chromatin architecture and functions leading up to ZGA.

## Materials and methods

### Whole embryo immunofluorescence

Eggs were fertilized and successfully dividing embryos were fixed in MAD (2 parts methanol, 2 parts acetone, 1 part DMSO) at 5-minute intervals at each stage when mitosis was likely to be occurring. After 1-3 hours of fixation, embryos were transferred to fresh MAD before storing at −20 °C for up to 3 months. Embryos were then gradually rehydrated into 0.5x SSC (75 mM NaCl, 7.5 mM sodium citrate, pH 7.0), bleached in 2% H_2_O_2_, 5% formamide and 0.5x SSC under direct light or 2-3 hours. Bleached embryos were blocked in 1x PBS, 0.1% Triton X-100, 2 mg/mL BSA, 10% goat serum, 5% DMSO for 16-24 hours at 4 °C. Primary antibodies were incubated at 2-10 μg/mL at 4 °C for 60 hours, washed in PBT (1x PBS, 0.1% Triton X-100, 2 mg/mL BSA) for 30 hours. Secondary antibodies were added at 2 μg/mL, covered from light and incubated for 60 hours at 4 °C before washing for 30 hours with PBT. Embryos were then gradually rehydrated into 100% methanol, stored overnight at −20 °C, then cleared with Murrays solution (2 parts benzyl benzoate, 1 part benzyl alcohol). Embryos were imaged with 20x or 40x objective on Zeiss LSM 780 confocal microscope. Once cells containing a mitotic spindle were identified, z-stacks were taken. Using Imaris, we performed 3D visualization and segmentation of mitotic spindles and metaphase plates to directly measure volumes. Cell size was measured in FIJI by manually tracing the cell in the z-stack where the spindle appeared the largest. We used this cell area measurement to calculate cell diameter. Interphase data were calculated using a published dataset (Jevtić and Levy, 2015), which used very similar methods to obtain measurements of nuclear size and cell size.

### Embryo extract preparation and sample reactions

#### Mitotic embryo extracts

Stage 3 and stage 8 mitotic embryo extracts were prepared as previously described (Wilbur and Heald, 2013). Briefly, eggs from at least 2 separate females were fertilized synchronously with ~1/2 of a fresh testes. After the appropriate amount of growth at 23 °C (1 hour and 45 minutes for stage 3 extract and 5.5 hours for stage 8 extracts), successfully dividing embryos were collected in 2 mL test tubes, washed with 5 times with XB (10 mM Hepes pH 7.8, 1 mM MgCl_2_, 0.1 mM CaCl_2_, 100 mM KCl, 50 mM sucrose) and 5 times with CSF-XB (10 mM Hepes pH 7.8, 2 mM MgCl_2_, 0.1 mM CaCl_2_, 100 mM KCl, 50 mM sucrose, 5 mM EGTA) containing protease inhibitors LPC (10 μg/mL leupeptin, pepstatin and chymostatin). Cytochalasin B (Cyto B) was added in the final wash for a final concentration of 20 μg/mL. Embryos were gently pelleted in a tabletop microcentrifuge at 1,000 rpm for 1 minute, then 2,000 rpm for 30 seconds. Embryos were then crushed in a swinging bucket rotor (Sorvall HB-6) at 10,200 rpm for 12 minutes at 16 °C. Cytoplasm was removed, placed on ice and immediately supplemented with 10 μg/mL LPC, 20 μg/mL CytoB, 1x energy mix (3.75 mM creatine phosphate, 0.5 mM ATP, 0.05 mM EGTA, 0.5 mM MgCl_2_), 0.25 μM cyclinBΔ90 and 5 μM UbcH10 C114S to induce metaphase arrest, and 0.6 μM rhodamine-labeled tubulin to visualize microtubules.

To examine endogenous mitotic chromosomes, 25-100 μL samples of embryo cytoplasm were incubated at 20 °C for ~1 hour or until spindles had formed. Samples were diluted 4-fold in CDB (250 mM sucrose, 10 mM Hepes pH 8.0, 0.5 mM EGTA, 200 mM KCl, 1 mM MgCl_2_) for 5-10 minutes, then diluted an additional 5-fold in CFB (5 mM Hepes pH 7.8, 0.1 mM EDTA, 100 mM NaCl, 2 mM KCl, 1 mM MgCl_2_, 2 mM CaCl_2_, 0.5% Triton X-100, 20% glycerol and 2% formaldehyde). Chromosomes were then layered on a 5 mL of CCB (5 mM Hepes pH 7.8, 0.1 mM EDTA, 100 mM NaCl, 2 mM KCl, 1 mM MgCl_2_, 2 mM CaCl_2_ 40% glycerol) and centrifuged onto coverslips at 5500 rpm (HB-6) for 20 minutes. Coverslips were removed and additionally fixed for 5 minutes in ice-cold methanol, and washed 5 times in 1x PBS, 0.1% NP-40 before moving on to immunostaining.

#### Interphasic embryo extracts

Embryos were first arrested in G2 using 150 μg/mL cycloheximide for 1.5 hours, then washed with ELB (250 mM sucrose, 50 mM KCl, 2.5 mM MgCl_2_, 10 mM Hepes pH 7.8) containing 10 μg/mL LPC, 200 μg/mL CytoB. Embryos were gently pelleted, then manually crushed with a pestle for 30 seconds before centrifuging at 10,000 g for 10 minutes. Cytoplasm was removed, placed on ice and immediately supplemented with 10 μg/mL LPC, 20 μg/mL CytoB, 1x energy mix, and 8% glycerol. Samples were aliquoted and flash frozen and stored at −80°C for up to 2 years. For immunofluorescence, embryo nuclei were thawed and directly fixed in ELB supplemented with 15% glycerol and 2.6% paraformaldehyde for 15 minutes with rocking at room temperature. Fixed nuclei were layered over a 5 mL cushion containing 100 mM KCl, 1 mM MgCl_2_, 100 μM CaCl_2_, 0.2 M sucrose, and 25% glycerol. Nuclei were spun onto coverslips at 1,000 g for 15 minutes at 16 °C. Coverslips additionally fixed and washed as described above.

#### Western blots

Stage 3 and stage 9 embryo extracts were prepared as described above and analyzed by Bradford to determine total protein concentrations. 25 μg protein was loaded per sample on 4-20% gradient gels (BioRad). Proteins were transferred overnight at 4 degrees onto nitrocellulose membranes, blocked in 5% milk in Tris Buffered Saline containing 0.1% Triton-X100 (TBST) for 1 hour at room temperature, then stained with primary antibodies for 1 hour at room temperature. After washing with PBST 5 times, blots were incubated with secondary antibodies containing infrared dyes for 1 hour at room temperature. After a final wash in PBST, blots were visualized on an LiCor Odyssey Imager.

### Egg extract preparation and sample reactions

Egg extracts from *X. laevis* were prepared as previously described (Maresca and Heald, 2006) For crude extracts, eggs were packed in a clinical centrifuge and crushed in an HB6 rotor for 16 minutes at 10,200 rpm. The cytoplasm was removed and supplemented with 10 μg/mL LPC, 20 μg/mL CytoB, 1x energy mix, and 0.6 μM rhodamine-labeled tubulin. For clarified extracts, crude extracts were centrifuged at 55,000 rpm for 2 hours, and then 30 minutes to pellet membranes (all steps at 4 °C). Supernatants containing soluble fraction of the cytoplasm were flash frozen and stored at −80 °C for up to 3 years.

#### Nuclei reactions and spin downs

Freshly prepared egg extracts were aliquoted into 20 μL reactions and sent into interphase with addition of 1x CA (0.4 mM CaCl_2_, 10 mM KCl, 0.1 mM MgCl_2_). After 5 minutes, sperm nuclei were added at 1000 nuclei/μL unless otherwise specified. Once nuclei had swollen, they were fixed and processed for immunofluorescence as stated above for embryo nuclei.

#### Mitotic chromosome reactions and spin downs

To form mitotic chromosomes from replicated sperm nuclei, purified sperm nuclei were added to 20 μL of interphase egg extract at 1000 nuclei/μL unless otherwise specified. After nuclei had swelled and chromatin was replicated (around 45 minutes), 30 μL fresh metaphase egg extract was added and spindles formed after ~45 minutes. To form mitotic chromosomes from stage 8 G2-arrested embryo nuclei, nuclei were first thawed on ice for 15 minutes before adding 1.5 mL of CSF-XB containing 10 μg/mL LPC. Nuclei were pelleted at 1600 g for 5 minutes at 4 °C. Nuclei pellets were resuspended in 10-15 μL of fresh CSF-XB (+LPC), and added at 1000 nuclei/μL to metaphase egg extracts, and mitotic spindles formed within 1 hour. Once successful formation of mitotic spindles and chromosome condensation was confirmed by taking a small sample and staining with Hoechst (1 μg/mL), samples were diluted 100-fold and fixed at the same time in ice-cold 1x XBE2 (5 mM Hepes pH 7.8, 100 mM KCl, 2 mM MgCl_2_, 0.1 mM CaCl_2_, 5 mM EGTA, 50 mM sucrose) containing 0.25% Triton X-100 and 2% formaldehyde. Fixed chromosomes were layered over a 5 mL cushion containing 1x XBE2 and 30% glycerol and spun onto coverslips at 5500 rpm for 20 minutes at 16 °C.

#### Mitotic spindle spin downs

To fix and spin down mitotic spindles instead of mitotic chromosomes, the same procedure was used except spindles were fixed in 1x BRB80 (80 mM PIPES pH 6.8, 1 mM MgCl_2_, 1 mM EGTA) containing 30% glycerol and 0,5% Triton X-100 and cushion buffer contained 1x BRB80 and 40% glycerol.

#### Anaphase reactions

Once mitotic spindles had formed, reactions were transferred to fresh tubes and 1x CA was added. After 40 minutes of interphase, an additional 0.5x CA was added to ensure full replication of DNA. Successful reactions were confirmed by staining a small sample with Hoechst. After 75 minutes of interphase, nuclei were swollen and an equal volume of fresh metaphase extracts were added to form a second round of metaphase spindles. Once formed, mitotic chromosomes were fixed and isolated using same procedures as described above.

#### Titration experiments

Anytime nuclei concentration in egg extracts was varied, the volume of extract fixed was also varied to keep the nuclei concentration per coverslip constant. We found this to be important for ensuring that any effects we observed were not due to titration of the antibody used for immunostaining.

#### Importin α experiments

These experiments were performed as described previously (Brownlee and Heald, 2019). Briefly, extracts were incubated with either DMSO or 10 μM palmostatin for 45 minutes at 20°C before using. Exogenous importin α (WT and NP) was purified from E. coli using previously published constructs and procedures (Brownlee and Heald, 2019).

### Immunofluorescence, imaging and analysis of chromosomes, spindles and nuclei from extracts

Once mitotic chromosomes, nuclei or spindles were fixed and isolated on coverslips, they were blocked overnight with 1x PBS, 3% BSA at 4 °C. Primary antibodies were added at 1μg/mL 2.5 μg/mL for 1 hour at room temperature and washed 5 times with 1x PBS, 0.1% NP-40. Secondary antibodies were added at 1 μg/mL for 1 hour, washed 5 times, then stained with Hoechst at 1 μg/mL for 10 minutes. Coverslips were washed two more times, then mounted using Vectashield without DAPI. Imaging was performed on an Olympus BX51 upright epifluorescence microscope using an Olympus PlanApo 60x oil objective for chromosomes and 40x air objective for nuclei and spindles. Images were analyzed in FIJI. Single chromosomes were manually selected and cropped from the rest of the image. Chromosome lengths were measured manually using the freehand line tool. Median intensity values were used to perform background subtractions in each channel and the abundance of a certain factor of interest was calculated by dividing the background-subtracted fluorescence intensity of the factor by the background-subtracted fluorescence intensity of Hoechst.

### Hi-C and contact probability analysis

#### Preparation of samples and sequencing

Hi-C was performed as previously described(Belaghzal et al., 2017). Mitotic chromosomes from either replicated sperm or stage 8 embryo nuclei were formed in 250 μL egg extract reactions containing 4000 nuclei/μL. Reactions were then diluted 48-fold in XBE2 containing 1% formaldehyde and 0.25% triton X-100. After 10 minutes of fixation with rocking at room temperature, samples were quenched for 5 minutes with 140 mM glycine before transferring to ice for 15 minutes. Chromatin was pelleted at 6,000 g for 20 minutes at 4°C, then resuspended in XBE2 containing 0.25% Triton X-100. Briefly, pallets were homogenized treated with 0.1% SDS (final concentration) and quenched with 1% Triton X-100 (final concentration) prior to overnight digestion with 400 U DpnII at 37 °C. Next day, enzyme was inactivated prior to biotin-fill with biotin-14-dATP for 4 hr at 23 °C. Subsequently, chromatin was ligated at 16 °C for 4 hours. After crosslinking was reversed by proteinase K at 65°C overnight. Sonicated ligation products were size selected for 100-350 bp products. Size selected products were end repaired followed by biotin-pull down with streptavidin. Prior to Illumina Truseq adapter ligation purified DNA fragments were A-tailed. PCR amplification and primer removal were last steps before final library was sequenced on Illumina HiSeq 4000 with PE50.

#### Hi-C analysis

Hi-C libraries processed by mapping to the *X. Laevis* 10 genome using the distiller pipeline (https://github.com/open2c/distiller-nf). Reads were aligned with bwa-mem, uniquely mapped reads were further processed after duplicate removal. Valid pair reads were binned at 1, 2, 5, 10, 25, 50, 100, 250, 500, and 1000kb in contact matrices in the cooler format (Abdennur and Mirny, 2019). Cooler files were normalized using Iternative balancing correction(Imakaev et al., 2012), excluding first two diagonals to avoid artifacts at short range.

#### Hi-C contact probability analysis

For contact probabilities balanced Hi-C data binned at 1kb was used to calculate contact frequency as function of genomic distance. From cooltools expected_cis, logbin_expected, and combined_binned_expected was used to generate average contact decay plots genome-wide (Venev et al. 2022). First, the contact frequency by distance for each chromosome was calculated using expected_cis. Data was grouped into log spaced bins with logbin_expected. Genome-wide average was calculated by combined_binned_expected.

---

**Figure 1, Supplement 1:**
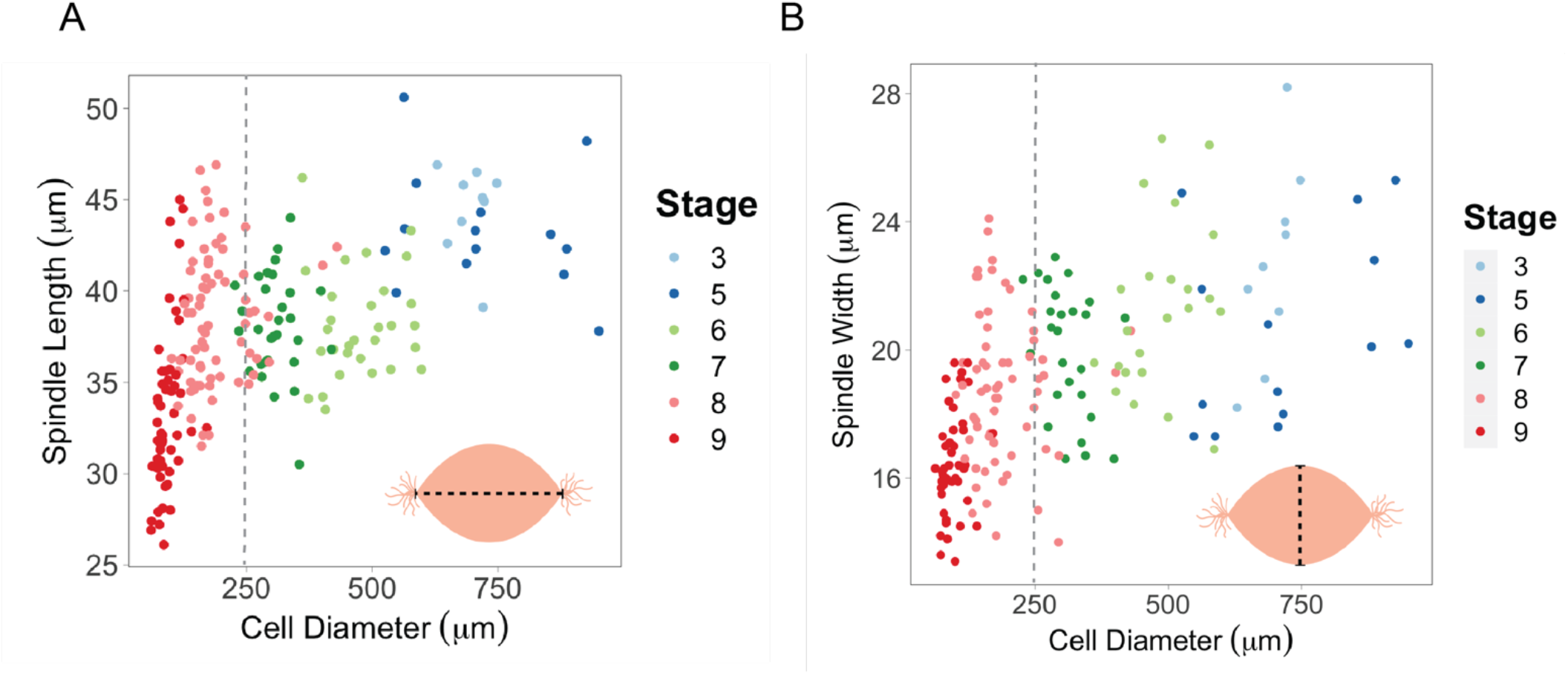
Spindle lengths or widths vs. cell size. (A) Spindle lengths and (B) widths plotted against cell diameter. Gray dotted lines show that spindle lengths plateau at around 250 μm while spindle widths continue to increase.

**Figure 1, Supplement 2:**
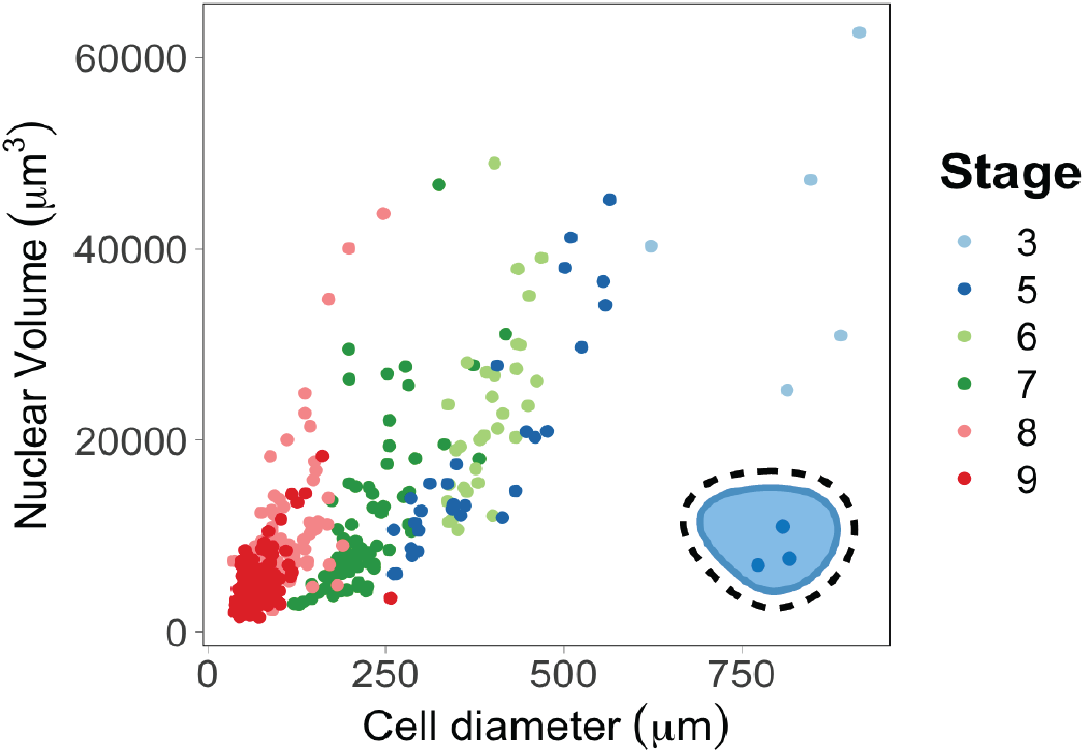
Nuclear volumes scale continuously with cell size. Nuclear volumes plotted against cell diameter. Raw data used with permission from Jevtić and Levy, *Current Biology* 2015.

**Figure 1, Supplement 3:**
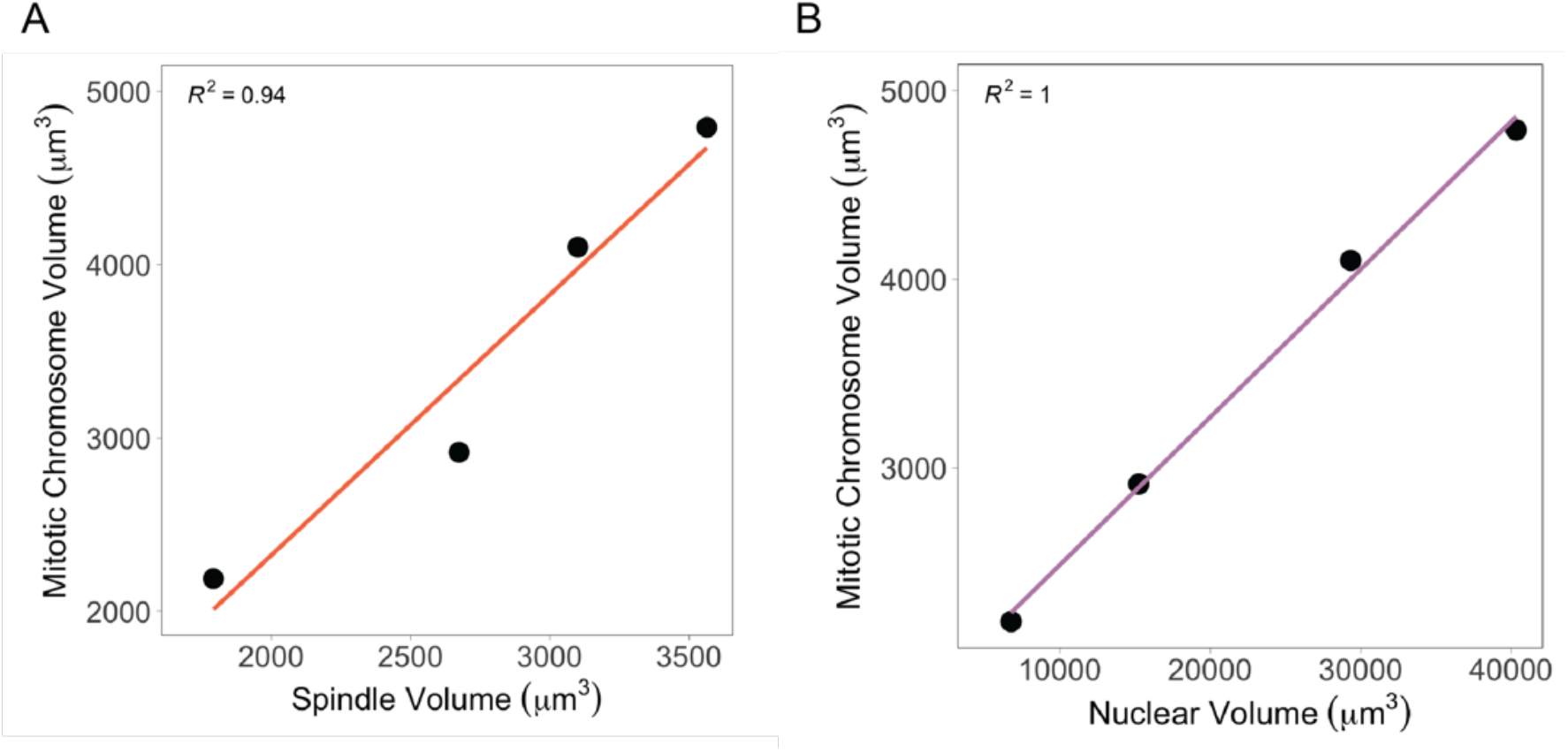
Mitotic chromosomes scale linearly with both nuclei and spindles. (A) Average mitotic chromosomes volumes plotted against (A) spindle volumes or (B) nuclear volumes, binned by cell diameter (bin 1 = 35.1 – 219 μm, bin 2 = 219 – 402 μm, bin 3 = 402 – 585 μm, bin 4 = 585-768 μm).

**Figure 1, Supplement 4:**
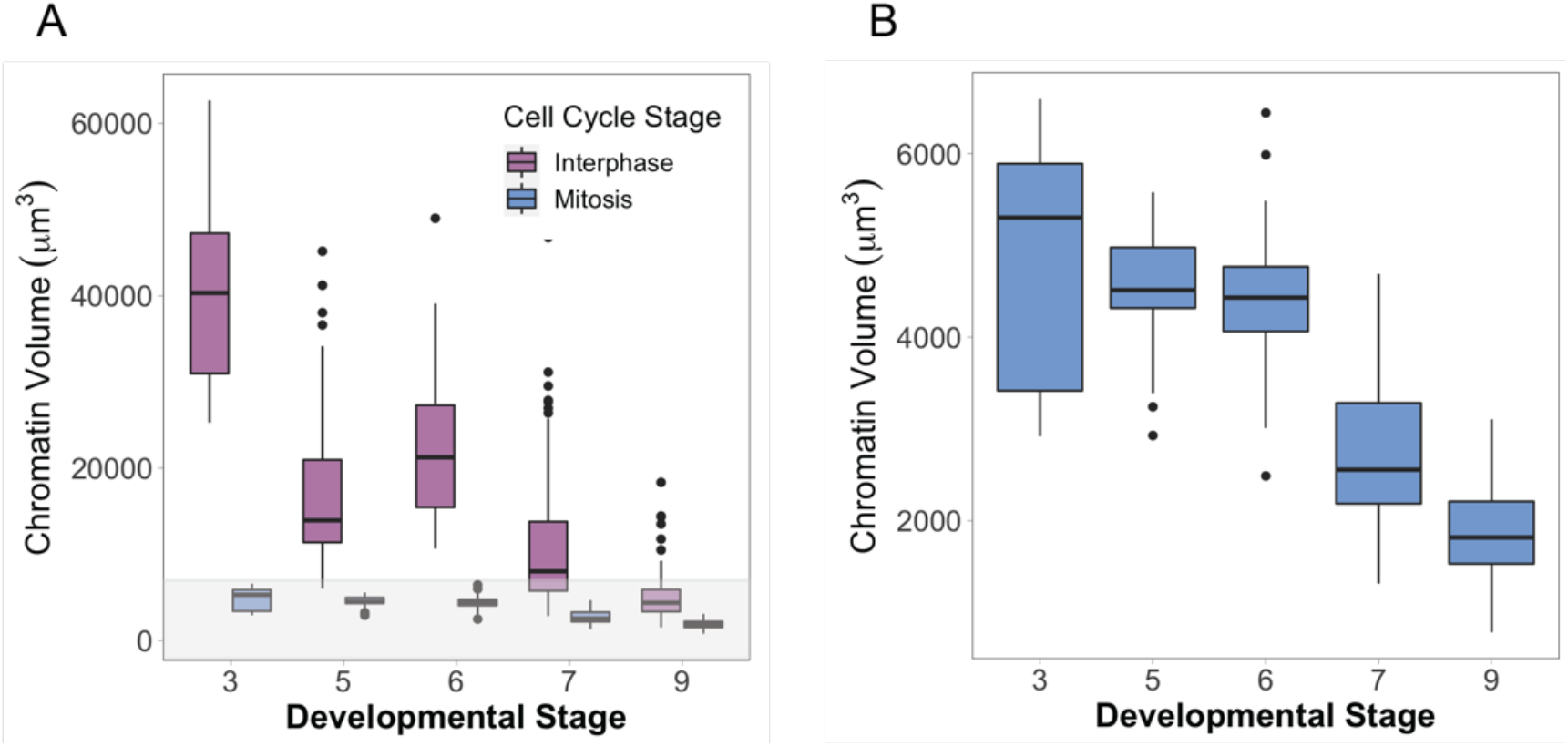
Nuclear vs. mitotic chromatin volumes during early cleavage divisions. (A) Chromatin volumes, either in interphase (purple) or metaphase (blue), binned by developmental stage. (B) Zoomed-in view of data shown in the gray panel in (A), only for mitotic chromosomes.

**Figure 3, Supplement 1:**
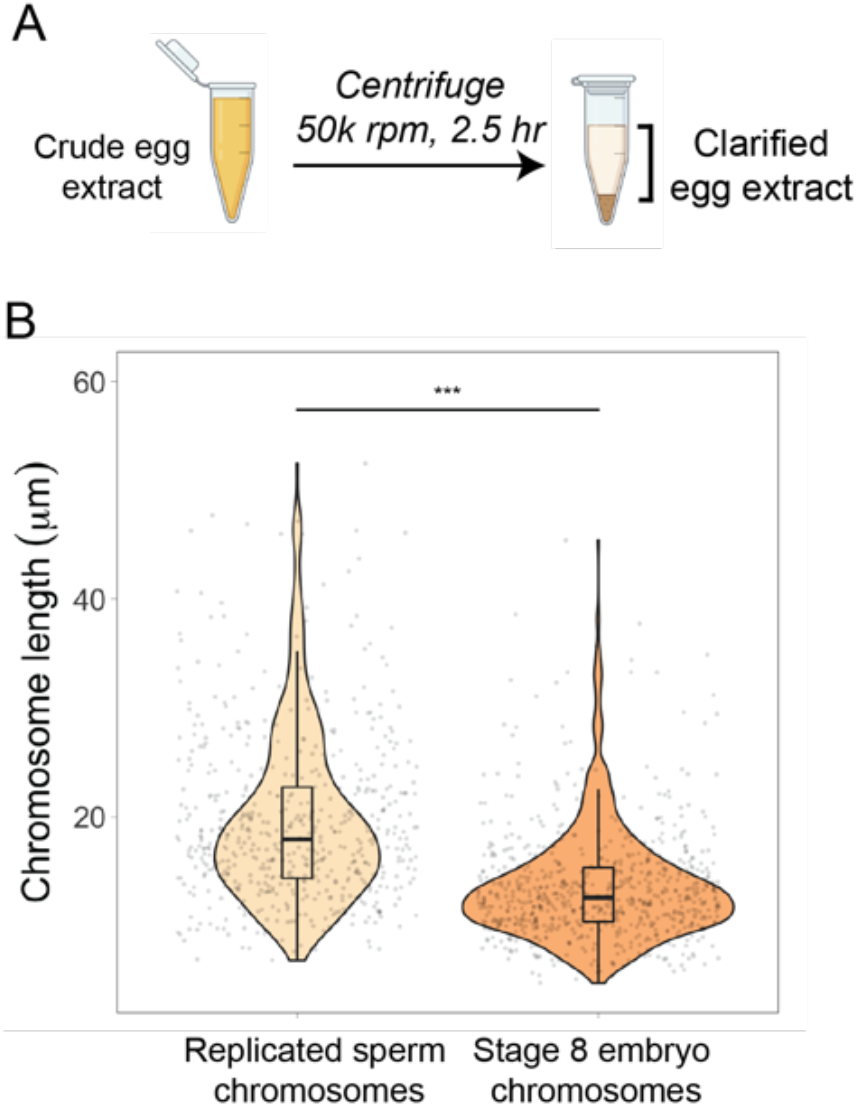
Clarified egg extracts recapitulate mitotic chromosome scaling. (A) Crude extracts were centrifuged at high speed to pellet membranes. (B) Quantification of mitotic chromosome lengths from either replicated sperm nuclei or stage 8 embryo nuclei added to clarified metaphase egg extracts. n=3 biological replicates, >50 chromosomes per replicate. *** denotes p <0.001 by one-way ANOVA statistical testing.

**Figure 3, Supplement 2:**
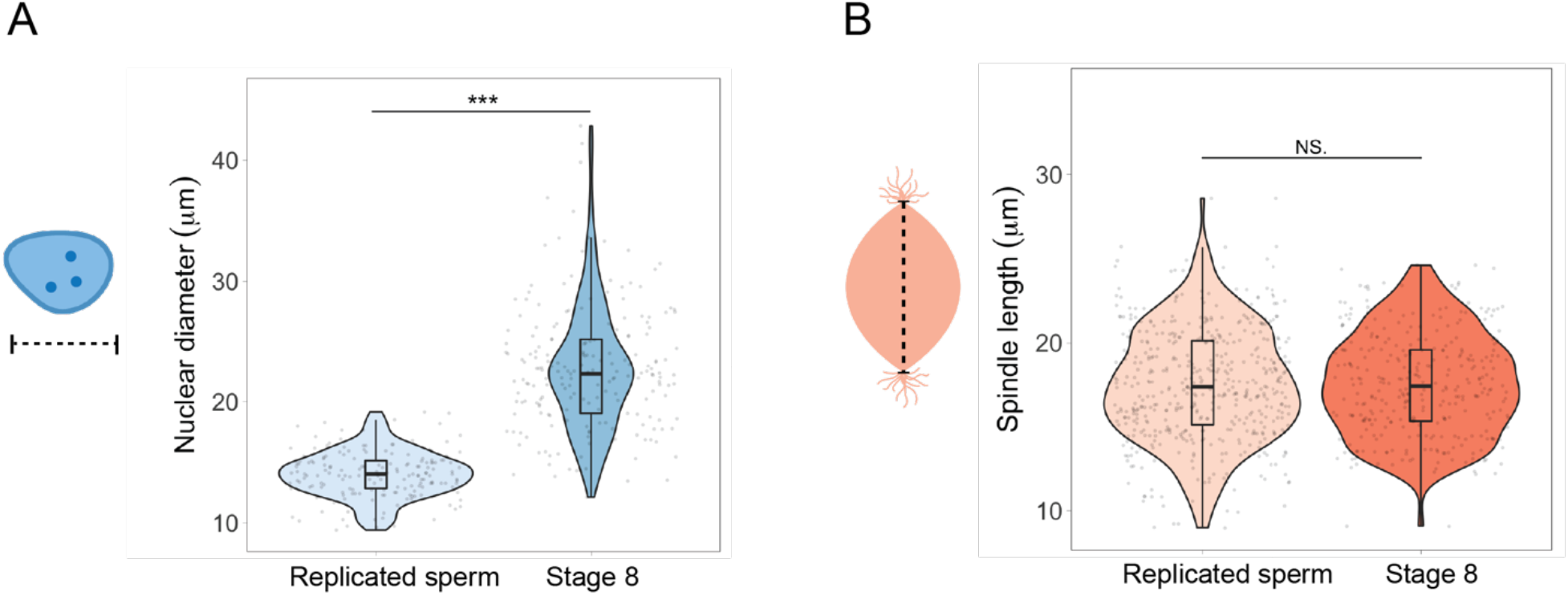
Nuclei and spindles do not scale with mitotic chromosome size in egg extracts. (A) Diameters of replicated sperm nuclei or stage 8 embryo nuclei just before placing into metaphase egg extracts. (B) Lengths of spindles formed around either replicated sperm nuclei or embryo nuclei in metaphase egg extracts. n=3 biological replicates, >50 structures per replicate. *** denotes p <0.001 by one-way ANOVA statistical testing.

**Figure 3, Supplement 3:**
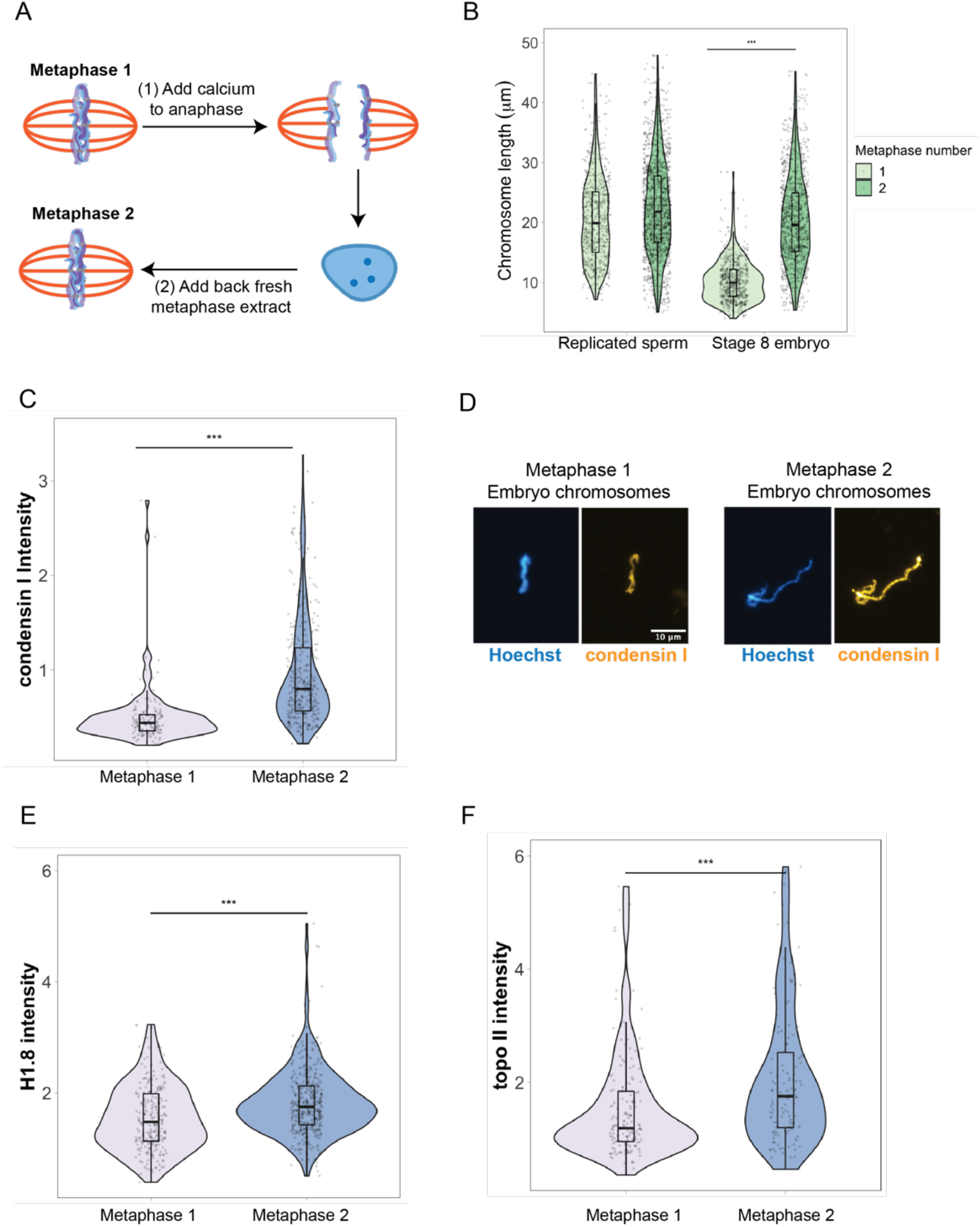
Scaling factors are re-loaded onto embryo chromosomes after an additional interphase in egg extracts. (A) Schematic of anaphase experiment. Calcium is added to send metaphase spindles containing embryo chromosomes into anaphase, and then interphase. After nuclei formed, fresh metaphase extract was added to trigger mitotic chromosome formation. (B) Quantification of chromosome lengths for the first and second metaphase comparing replicated sperm and stage 8 embryo mitotic chromosomes. (C) Abundance of condensin I on stage 8 embryo chromosomes in the first or second metaphase. (D) Representative images of embryo chromosomes from metaphase 1 or metaphase 2, stained for condensin I. (E-F) Abundance of H1.8 and topo II on stage 8 embryo chromosomes in the first or second metaphase. Based on median values, condensin I increased 2-fold, H1.8 increased 1.2-fold and topo II increased 1.5 fold from the first to second metaphase. n=3 biological replicates, >50 chromosomes per replicate. *** denotes p <0.001 by one-way ANOVA statistical testing.

**Figure 5, Supplement 1:**
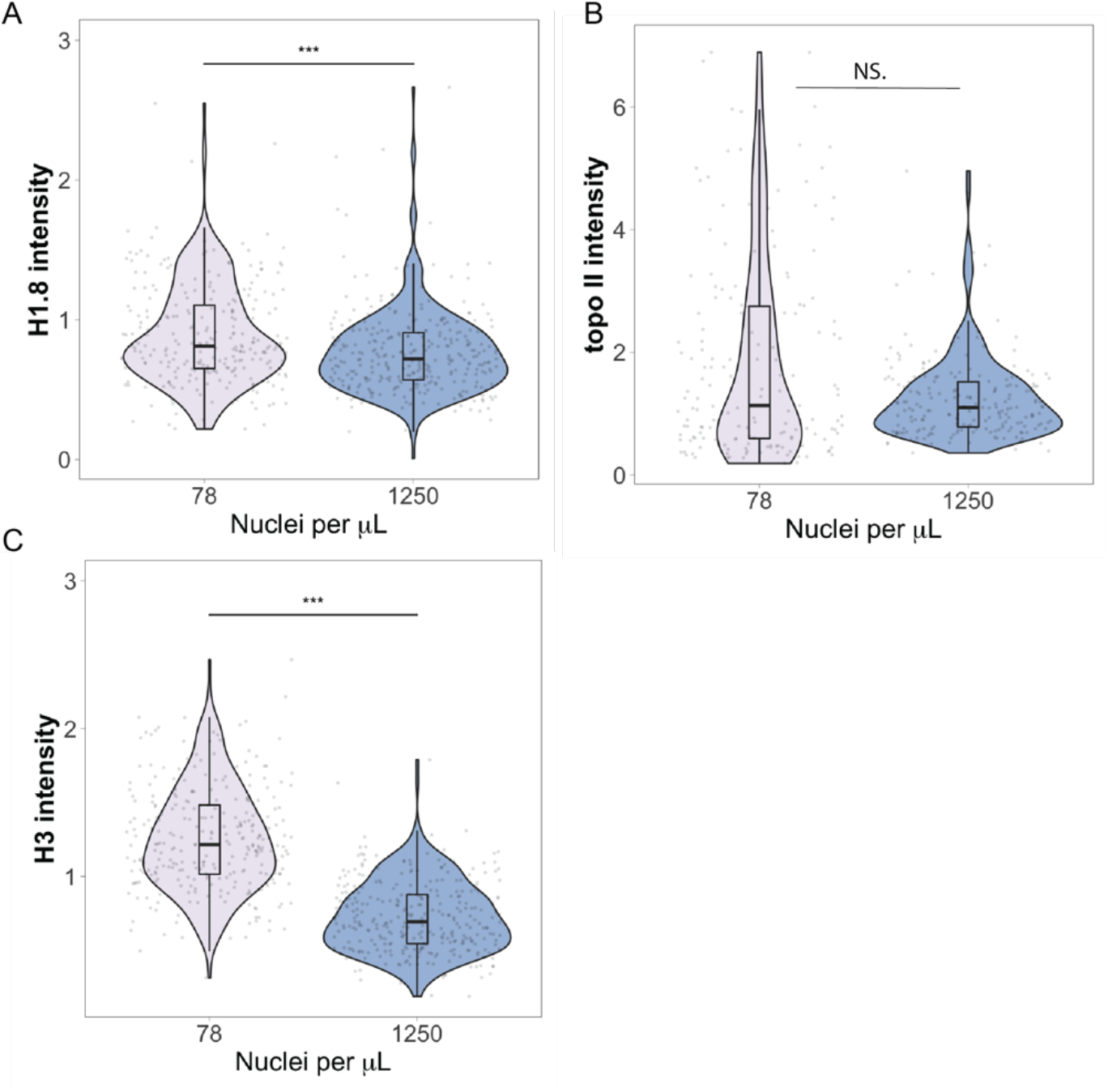
topo II and H1.8 exhibit less of a titration effect than condensin I. Normalized fluorescence intensity of (A) H1.8, (B) topo II, and (C) H3 on sperm mitotic chromosomes in samples containing low or high concentrations of nuclei. n=3 biological replicates, >50 chromosomes per replicate, *** denotes p <0.001 by one-way ANOVA statistical testing.

**Figure 5, Supplement 2:**
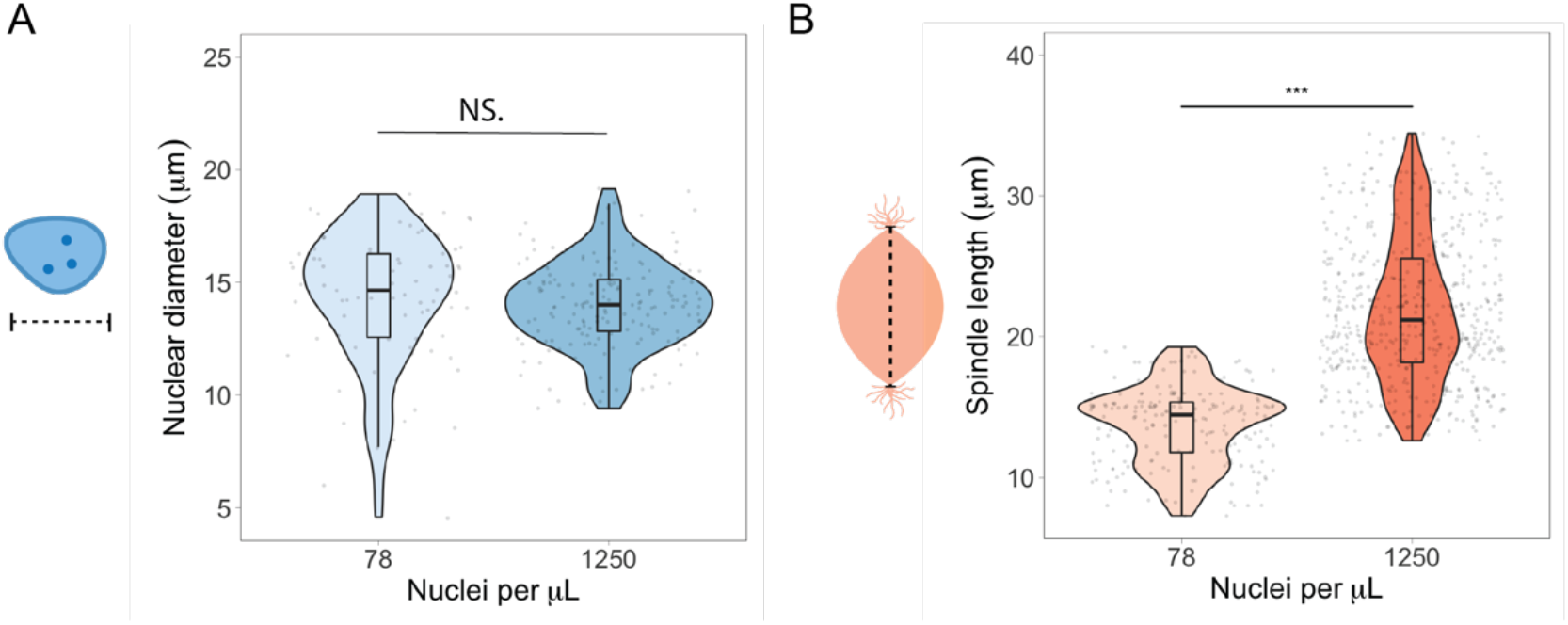
N/C ratio does not regulate spindle or nuclear size. (A) Nuclear diameters plotted for samples containing low or high concentrations of sperm nuclei. (B) Spindle lengths plotted for samples containing low or high concentrations of sperm nuclei. n=3 biological replicates, >50 structures per replicate, *** denotes p <0.001 by one-way ANOVA statistical testing.

**Figure 5, Supplement 3:**
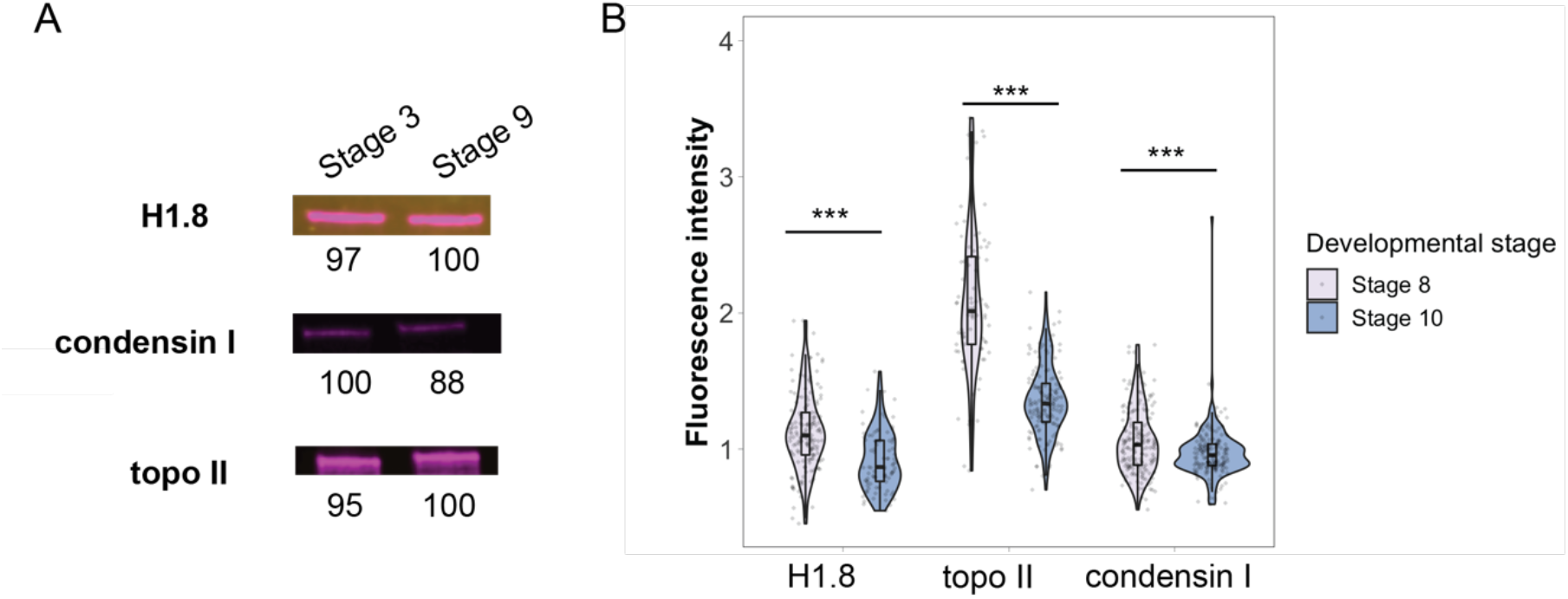
Titration of nuclear factors during embryogenesis. (A) Western blots of whole embryo extracts from early (stage 3) or late (stage 9) blastula stages. Numbers below each band indicate the relative differences in signal intensity, normalized to the highest intensity band for that antibody. (A) Immunofluorescence of nuclei from stage 8 or 10 embryos showing the depletion of nuclear factors in later stages. n=3 biological replicates, >50 chromosomes per replicate, and *** denotes p <0.001 by one-way ANOVA statistical testing.

**Figure 5, Supplement 4:**
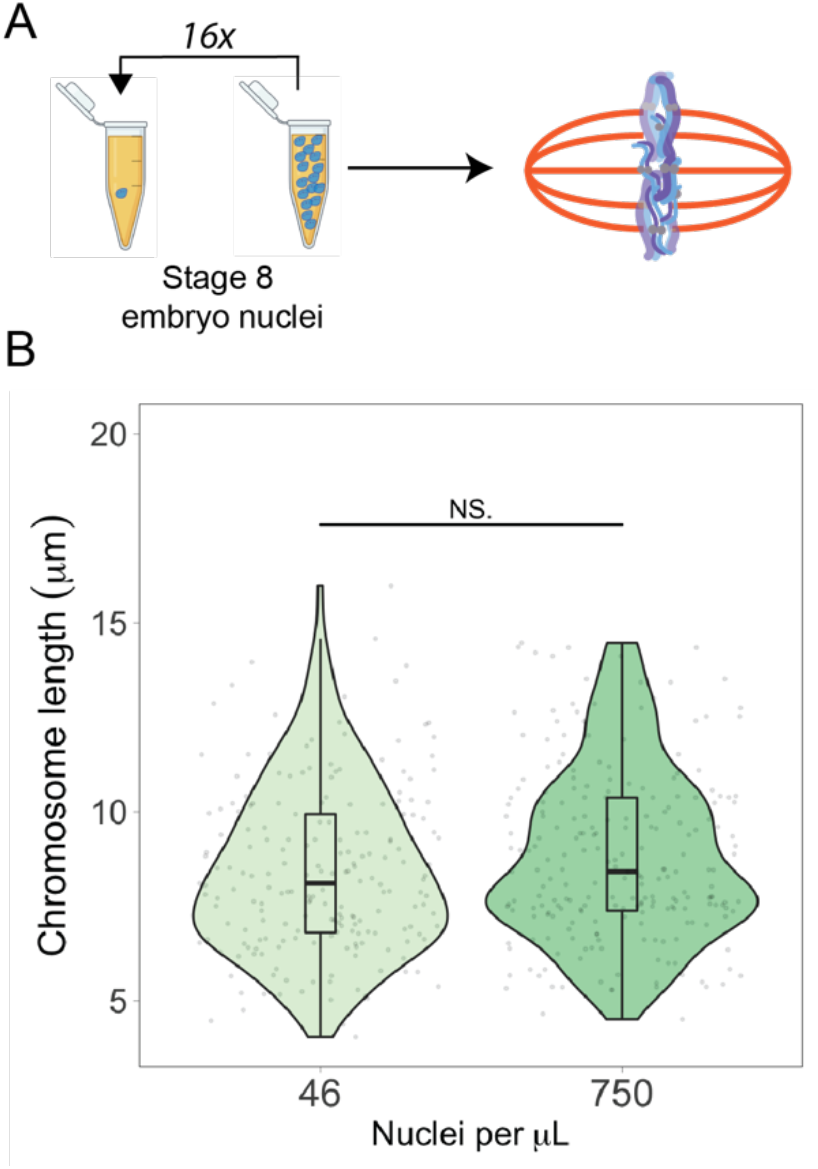
Embryo mitotic chromosome lengths are not affected by nuclei density. (A) Schematic of experiment. Stage 8 embryo nuclei were added to metaphase egg extracts at N/C ratios spanning a 16-fold range. (B) Quantification of mitotic chromosome lengths in samples containing high or low concentrations of stage 8 embryo nuclei. n=3 biological replicates, >50 chromosomes per replicate, and *** denotes p <0.001 by one-way ANOVA statistical testing.

## Acknowledgements

We would like to thank Yoshiaki Azuma (University of Kansas) and Susannah Rankin (OMRF) for their generous donation of antibodies against topo II and x-CAPG. We would like to thank J. Smolka, H. Cantwell, G. Cavin-Meza, X. Liu and S. Coyle for their feedback on the manuscript. Some illustrations were made with the help of Rebecca Konte and Biorender.com. This work was supported by an R35 from NIH NIGMS to R.H. and a Jane Coffin Childs Memorial Fund fellowship to C.Y.Z.

